# Metabolic responses of normal rat kidneys to a high salt intake

**DOI:** 10.1101/2023.01.18.524636

**Authors:** Satoshi Shimada, Brian R. Hoffmann, Chun Yang, Theresa Kurth, Andrew S. Greene, Mingyu Liang, Ranjan K. Dash, Allen W. Cowley

## Abstract

In the present study, novel methods were developed which allowed continuous (24/7) measurement of blood pressure (BP) and renal blood flow (RBF) in freely moving rats and the intermittent collection of arterial and renal venous blood to estimate kidney metabolic fluxes of O_2_ and metabolites. The study determined the effects of a high salt (HS) diet upon whole kidney O_2_ consumption and the metabolomic profiles of normal Sprague Dawley (SD) rats. A separate group of rats was studied to determine changes in the cortex (Cx) and outer medulla (OM) tissue metabolomic and mRNAseq profiles before and following the switch from a 0.4% to a 4.0% NaCl diet. Significant changes in the metabolomic and transcriptomic profiles occurred with feeding of the HS diet. A progressive increase of kidney O_2_ consumption was found despite a reduction in expression of most of the mRNA encoding enzymes of TCA cycle. Increased glycolysis was evident with the elevation of mRNA expression encoding key glycolytic enzymes and release of pyruvate and lactate from the kidney in the renal venous blood. Glycolytic production of NADH is used in either the production of lactate or oxidized via the malate aspartate shuttle. Aerobic glycolysis (e.g., Warburg-effect) may account for the needed increase in cellular energy. The study provides evidence that kidney metabolism responds to a HS diet enabling enhanced energy production while protecting from oxidate stress and injury.

## Introduction

The process of converting daily food intake into energy and biomass is a complex process with some operating relatively inefficiently as essential responses for short term survival while others operating in more efficient and sustained ways required in face of sustained chronic stressors^1-3^. Malnourishment is one such chronic stressor, but industrialized societies are largely faced with dietary excesses and trying to deal with the consequences of an overload of nutrients and related condiments. Among the most onerous of these condiments is salt (NaCl) which is no longer needed for food preservation but is consumed in amounts greatly in excess of what is needed for survival of the organism. These excesses of dietary salt increase the risk of hypertension especially in salt-sensitive individuals. Nearly half of hypertensive patients are blood pressure (BP) salt-sensitive^4-6^ and exhibit a 3-fold greater risk of chronic kidney disease (CKD)^7-9^. Yet, many of the underlying mechanisms remain poorly understood.

Many advances have been made in the understanding of the neural and endocrine controllers of body metabolism^10, 11^. Studies of whole kidney and segmental nephron renal oxygen and substance metabolism under normal conditions in vitro and in vivo have been widely conducted and are summarized in textbooks^12, 13^. The effects of what we eat (dietary content) upon these metabolic responses and energy production within vital organs have recently received increased attention driven by the obesity epidemic and associated diabetes in industrialized nations. However, because of the technical difficulties of obtaining repeated samples from the same individuals, only a few studies have examined the effects of dietary salt on kidney metabolism which is the subject of the current study. Kidneys have one of the highest specific metabolic rates among all organs estimated in humans to be over 400 kcal/kg tissue/day, which is the same as the heart, twice as high as the liver and the brain, and much higher than other organs^13, 14^. Kidney metabolism is tightly linked to renal tubular transport activities which are critical for the regulations of fluid and electrolyte homeostasis and blood pressure.

Traditionally, studies of organ metabolism have been limited to examination of a relatively few metabolites^15, 16^. With the emergence of large-scale mass spectrometry and analysis tools (aka metabolomics), it is currently possible to identify and prioritize several thousands of detected features providing a comprehensive analysis in a tissue specimen of the changes in metabolism that occur across various organs of the body. The potential power of these methods was demonstrated recently in a study by Jang et al.^17^ who analyzed the arterial and venous blood of 11 organs in fasted pigs and mapped more than 700 cases of organ-specific metabolite production or consumption. More relevant to the current study, Rinschen et al.^18^ recently carried out a metabolomic and proteomic analysis of the kidney glomeruli and cortical tissue to determine the effects of a high salt diet in a naturally occurring rat model of salt-sensitive hypertension, the Dahl SS rat. As yet, no one has characterized in the absence of anesthesia and surgical stress how kidneys in either normal or diseased subjects or rodent may be metabolically altered by a high salt diet.

Compared to genomics or transcriptomics, the field of metabolomics is yet in an infant stage. The metabolome by definition represents the complete set of metabolites found in a biological sample and build upon the genetic blueprint of an organism. As the end product of gene expression it represents a sensitive method to measure biological phenotypes and in contrast to measurements of gene expression can change rapidly in the timescale of seconds or minutes thereby reflecting the function of the tissue at a given time point. Metabolomics detects the products of various stages of metabolism which have a wide range of functions and include amino acids, alcohols, vitamins, polyols, organic acids, and many other types of small molecules generally smaller than 1500 MW in size. Integrating transcriptomics-metabolomics dataset reveals the bidirectional and multi-faceted interactions between DNA and RNA elements that lead to observable phenotypes and provides insights into what is going on in a biological system.

Mass spectrometry can measure molecules with wide ranging physical properties which may vary in polarity and from highly water-soluble organic acids to very nonpolar lipids^19, 20^. Metabolomic technology platforms must therefore divide the metabolome into subsets of compounds based on polarity, common functional groups, or structural similarity. Varying methods of sample preparation are required to optimize the analysis of each class of compounds. Frequent refinements of these methods are common in this rapidly evolving field, the precision of instruments used for analysis differs and the degree of certainty in metabolite identification varies between laboratories.

In the present study, we have developed a novel system which allows us to measure blood pressure (BP) and renal blood flow (RBF) continuously (24/7) in freely moving rats, and collect arterial and renal venous blood repeatedly. Normal Sprague-Dawley (SD) rats were studies to determine global metabolic fluxes and O_2_ consumption determined by differences between the arterial input and the two major exit routes, the renal venous blood and urinary output. Rats were studied at weekly intervals following an increase of dietary salt intake from 0.4% (LS) to 4.0% (HS) NaCl. The effects of a HS diet upon the transcriptomic and metabolomic profiles of the kidney cortex and outer medulla were also determined from groups of SD rats in which kidneys were removed at days 14 and 21 of the HS diet and compared to low salt fed SD rats. Novel analytical approaches for tissue, plasma and urinary metabolism were developed to determine metabolic profiles by 4 modes (C18+/- and HILIC+/-) using a Thermo Q-Exactive Orbitrap coupled to a dual-channel Vanquish Ultra-Performance Liquid Chromatography (UPLC) system. Tissue, plasma, and urinary samples were analyzed with separation over a hydrophobic (C18) and hydrophilic (HILIC) column under both positive and negative polarity to maximize identification of metabolites^21^. Differential metabolites include numerous lipids, amino acids, and bioenergetic compounds, among others. The results of the study show that even normal SD rats undergo enormous shifts in transcriptomic and metabolomic profiles in response to eating a HS diet. These adaptations appear necessary to sustain vital physiological functions of the kidney and to prevent injury in face a great increase of the metabolic workload placed upon the kidney when subjected to sustained high salt diets.

## Methods

### Animals

Male SD rats were purchased from Envigo (Indianapolis, IN) and housed in environmentally controlled rooms with a 12-h light/dark cycle. Rats had free access to 0.4% NaCl AIN-76A diet (LS) (Dyets, Bethleham, PA) and water ad libitum. All protocols were approved by the Medical College of Wisconsin Institutional Animal Care and Use Committee.

### Chronic RBF measurement and blood sample collection

Rats (n=7, 10-11 weeks of age) were performed renal blood flow (RBF) probe (Transonic, Ithaca, NY) implantation and femoral arterial catheterization as previously described^22, 23^. Briefly, rats were anesthetized with isoflurane and arterial catheter was inserted as previously described^24-26^. Following an abdominal incision, RBF probe was implanted on left renal artery and the cable was exposed at nape of the neck via the subcutaneous route. In addition to the RBF probe implantation, renal venous catheter (MRE025, BRAINTREE, MA) was inserted through the femoral vein and placed in the left renal vein and secured to the luminal wall with 10-0 nylon (**Figure S1**). Three percent heparinized saline was infused at a rate of 100 μL/h through arterial and renal venous catheter throughout the study. RBF and blood pressure (BP) via arterial line were measured by conscious freely moving rats and recorded on average of every minute for 24 h/day. After 7-10 days of recovery period, 200 μL of arterial and renal venous blood were sampled and that blood was replace from donor rats before and following 7, 14 and 21 days after the switch in diet from 0.4% (LS) to 4.0% (HS) salt diet (Dyets Inc, Bethlehem, PA). Blood gases (pO_2_ and pCO_2_; mmHg), electrolytes, total hemoglobin concentration (Hb; g/dL), and oxyhemoglobin saturation (SHbO_2_; %) were immediately measured by radiometer (ABL800 FLEX, Brea, CA). Overnight urine (18 hours) from the day before the blood draw were collected on ice. The kidneys were collected either at 14 days of HS (HS14) or 21 days of HS (HS21). The kidneys of only LS fed SD rats were also collected for comparison. The collected kidneys (n=5 for each group for metabolomics and mRNAseq analysis) were dissected to cortex and outer medulla and snap frozen with liquid nitrogen. Plasma, urine and tissue were stored in -80°C until further analysis. RBF is often normalized by kidney weight, but it is impossible to repeatedly measure the kidney weight of the same rats and even measuring body weight repeatedly is difficult in this model. As salt did not alter the kidney weight of surgical sham control rats (n=5 for LS group, n=6 for HS group) (**Table S1**), data normalization to body weight was performed.

Since glomerular filtration rate (GFR) experiments and blood draws were performed during the daytime, the average RBF over a 12-hour period during the daytime (6AM-6PM) was used for the following calculations.

### Chronic GFR measurement

GFR was measured by separate group of rats (n=6) by transcutaneous measurement of FITC-sinistrin as previously described^22, 27, 28^. Briefly, an indwelling inferior vena cava catheter was implanted 7-10 days before GFR measurement via femoral vein. An abdominal median incision was performed to be considered as a surgical sham of the other group. GFR was measured before and following 7, 14 and 21 days after the switch in diet from LS to HS. GFR (mL/min/100 g body wt) was defined as 21.33 mL/100 g body wt, the conversion factor calculated by Friedemann et al.^29^, divided by FITC-sinistrin half-life (min). From the GFR and RBF from the other group, filtration fraction was calculated by the formula below:

*Filtration fraction = GFR / (2 x RBF x (1-Hct))*, where Hct is the hematocrit.

Tubular reabsorption of sodium was estimated as below^30^:

*GFR x whole blood Na*^*+*^ *(measured in RBF group rats by radiometer) – Urine flow x urinary Na*^*+*^ *(measured in RBF group rats by radiometer)*.

### Metabolomics analysis

*Plasma/Urine Metabolite Extraction*. Metabolites were extracted from 20 μL of plasma and 20 μL of urine from each SD rat in the study according to standard operating procedures in the Mass Spectrometry and Protein Chemistry Service at The Jackson Laboratory^21^. Metabolites were extracted using 500 μL of an ice cold 2:2:1 methanol:acetonitrile:water (MeOH:ACN:H_2_O) buffer; the sample was part of the water fraction. Caffeine, 1-napthylamine, and 9-anthracene carboxylic acid were all added at 0.5 ng/ μL in the extraction buffer as internal standards. Each sample was then vortexed for 30 seconds on the highest setting, subject to one minute of mixing with the Tissue Lyser II in pre-chilled cassettes, and then sonicated at 30 Hz for 5 minutes of 30 seconds on 30 seconds off in an ice water bath. Samples were then placed in the -20°C freezer overnight (16 hours) for extraction. Following the extraction, samples were centrifuged at 21,000 x g at 4°C and supernatant from each metabolite extract was equally divided into five 2 mL microcentrifuge tubes. Each sample supernatant was divided into five equal volume aliquots, one for each of the four modes and the rest to create equal representation pools of all samples, one for each mode. Each aliquot was then dried down using a vacuum centrifuge for storage at -80°C until further use.

#### Tissue Metabolite Extraction

Metabolites were extracted from 20 mg of kidney cortex and medulla from each SD rat in the study according to standard operating procedures in the Mass Spectrometry and Protein Chemistry Service at The Jackson Laboratory^21^ as described for the plasma and urine samples with slight modification. Metabolites were extracted using 1000 μL of an ice cold 2:2:1 methanol:acetonitrile:water (MeOH:ACN:H_2_O) buffer containing internal standards as above per 20 mg of sample to ensure the extraction equivalents were normalized. Each sample had a 5 mm stainless steel bead added, then were pulverized in extraction buffer for two minutes usingTissue Lyser II. Samples were then placed in the -20°C freezer overnight (16 hours) for extraction and the supernatant was collected as with the urine/plasma samples. Each sample supernatant was divided into five equal volume aliquots, one for each of the four modes and the rest to create equal representation pools of all samples, one for each mode. Each aliquot was then dried down using a vacuum centrifuge for storage at -80°C until further use.

#### Discovery Metabolomics Analysis

The Mass Spectrometry and Protein Chemistry Service at The Jackson Laboratory performed four mode metabolomics analysis using a Thermo Q-Exactive Orbitrap mass spectrometer coupled to a dual-channel Vanquish Ultra-Performance Liquid Chromatography system as described previously^21^. All samples were subject to the four modes of analysis consisting of a 25-minute gradient over a hydrophobic C18 column (Agilent InfinityLab Poroshell 120 EC-C18, #699775-902T) and a hydrophilic HILIC column (Agilent InfinityLab Poroshell 120 HILIC-Z, #689775-924) column in positive and negative polarity. The C18 runs used a gradient from 99.8% H_2_O with 0.2% acetic acid (Solvent A1) to 99.8% ACN with 0.2% acetic acid (Solvent B1). The specific C18 gradient consisted of the following steps: 0-1 minutes at 98% A1/2% B1, 1-13 minutes from 98% A1/2% B1 to 10% A1/90% B1, 13-15 minutes at 10% A1/90% B1, 15-16 minutes from 10% A1/90% B1 to 98% A1/2% B1, and was re-equilibrated from 16-25 minutes at 98% A1/2% B1. All HILIC positive runs used a gradient from 10 mM ammonium formate in H_2_O with 0.1% formic acid (Solvent A2) to 90% ACN with 10 mM ammonium formate in H_2_O with 0.1% formic acid (Solvent B2). HILIC negative runs utilized a gradient of 10 mM ammonium acetate in H_2_O, pH 9.0 with 0.1% AffinityLab Deactivator Inhibitor (Agilent, #5191-3940; Solvent A3) to 85% ACN with 10 mM ammonium acetate in H_2_O with 0.1% AffinityLab Deactivator Inhibitor (Solvent B3). The 25 minute gradient for the HILIC modes consisted of the following steps (A/B refer to A2/A3 B2/B3 for the respective HILIC mode): 0-1 minutes at 2% A/98% B, 1-11 minutes from 2% A/98% B to 30% A/70% B, 11-12 minutes from 30% A/70% B to 40% A/60% B, 12-16 minutes from 40% A/60% B to 95% A/5% B, was held at 95% A/5% B from 16-18 minutes, 18-20 minutes from 95% A/5% B to 2% A/98% B, and was re-equilibrated from 20-25 minutes at 2% A/98% B.

Each sample was reconstituted in 25 μL of 95% H_2_O/5% ACN for C18 modes and 95% ACN/5% H_2_O for HILIC modes. The sample run sequence was randomized (Random.org) and two technical replicates for each sample were injected at 10 μL (represents ∼2 μL of fluid samples or 2 mg of tissue sample starting volume weight, respectively, per run). Quality control pooled samples representing all samples within the specific fluid/tissue were run at the beginning and end of the run set at concentrations equivalent to the samples. These pooled samples were used for normalization through a quality control batch correction of the runs over time to account for technical variance. All instrument settings were set as described in a previous study^21^.

#### Metabolomics Data Analysis

The RAW data files were analyzed using Thermo Compound Discoverer (v3.2.0.421) according to supplementary methods from a previous study at The Jackson Laboratory^21^. Spectra in the data were subject to a blank background subtraction to remove contaminant peaks (S/N threshold = 2). Additionally, all data was subject to a quality control correction selecting for peaks only consistently detected in the pool for normalization. The MS1 and MS2 data was searched against the Thermo mzCloud database, ChemSpider database, Metabolika Pathways, and mzLogic predicted composition in the Compound Discoverer workflow. All data was then filtered for consistency of detection using a coefficient of variation ≤ 35% in any group, as well as for quality of MS2 spectral matching using an MS2 FISH coverage filter > 10 in Compound Discoverer. Differential comparisons were performed comparing normalized abundances and p-values were calculated using the Tukey HSD test (posthoc) after an ANOVA test. From there the p-values were adjusted for stringency in the multiple testing using the Benajmini-Hockberg algorithm. Further analysis was performed using a combination of Compound Discoverer, custom R analysis, and MetaboAnalyst as needed.

### Lactate assay

Tissue and plasma lactate concentration was validated by a commercially available kit according to the manufacturer’s instructions (PicoProbe™ Lactate Fluorometric Assay Kit, Bio Vision Cat# K638-100).

### RNAseq analysis

RNAseq analysis and data analysis were performed at Novogene (Durham, NC). Methods are described in a separate document.

### Determination of kidney O_2_ and metabolites extraction ratio

Whole blood O_2_ extraction was calculated as previously described^31^:

*Oxygen content (mL/dL) = (1*.*31 x Hb (g/dL) x SHbO*_*2*_*) + (0*.*003 x pO*_*2*_)

*Oxygen delivery (DO*_*2*_*) (mL/min) = RBF (daytime 12 hr; mL/min) x arterial oxygen content (mL/mL)*

*Oxygen consumption (VO*_*2*_*) (mL/min) = RBF (daytime 12 hr) x arterial-renal venous content difference*

*Oxygen extraction ratio = VO*_*2*_*/DO*_*2*_

Plasma metabolites flux was calculated similarly.

*Metabolites flux = (RBF x (1-Hct) x arterial-renal venous metabolites difference – Urine flow x urinary metabolites)/ (RBF x (1-Hct) x arterial metabolites)*

#### Data analysis of metabolomics

Compound names were converted to the corresponding names available in the Metaboanalyst 5.0 database (https://www.metaboanalyst.ca) (analyzed September-December 2022) by “Compound ID Conversion” and those compounds are used as “reference metabolome” which can be detected based on our analytical platform. Sparse partial least squares discriminant analysis (sPLS-DA) for all named compounds in each of the 4 modes (C18+/- and HILIC+/-) was performed. The number of components was fixed at 5 and variables per component at 20. Heat map with the hierarchical clustering analysis was performed for significantly changed compounds between LS, HS7, HS14 and HS21 (ANOVA Fisher’s LSD p<0.05) with a Euclidean distance measure and by the Ward algorithm. Enrichment analysis for tissue metabolites was performed for significant differences by t-test (p<0.05) in each of HS14 and HS21 compared to LS. Enrichment analysis for plasma metabolites was performed for significant different metabolites by linear models with covariate adjustments (p<0.05) in arterial and venous differences. Those data analyses of metabolomics were performed using Metaboanalyst^32^.

### Statistical analysis

Continuous values are presented as the means ± standard error of the means (SEM). Statistical comparisons were made using a t-test for two-group comparisons, and analysis of variance (ANOVA) followed by Holm Sidak’s post-hoc test for multiple between-group comparisons. A p<0.05 was considered significant. ROUT test was performed for outlier test (Q=5%). The error in the values obtained by combining values in tubular reabsorption of Na^+^ calculation with errors was indirectly estimated based on the error propagation formula shown below (*M*: Mean, *e*: error)^33^:

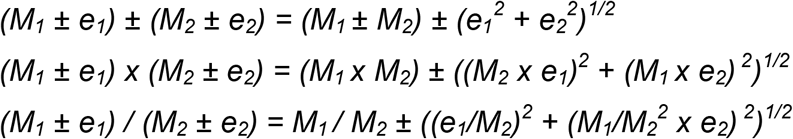

## Results

### Effects of high salt diet on arterial pressure, renal blood flow, and O_2_ extraction

Average 24 hr mean arterial pressure (MAP) of SD rats slightly but significantly increased from 114 ± 2 to 119 ± 2 mmHg (p<0.05) in 3 days after switching the diet from LS to HS and maintained at that level throughout the study (**Figure 1A**). Average 24 hr RBF rose nearly 20% during the first 3 days from 10.0 ± 0.6 to 11.6 ± 0.6 mL/min (p<0.05; **Figure 1B**) and was sustained at this elevated level throughout the 21 days of the HS diet. Normalized by the body weight changes determined in the surgical sham control rats (**Table S2**), the increase in RBF with the HS diet remained statistically significant at HS7, HS14 and HS21 (**Table S3**). Renal vascular resistance (RVR) did not change by HS (p=0.75). An example of a SD rat in which arterial pressure and RBF were recorded continuously (24/7) before and 21 days following the switch to the HS diet is illustrated in **Figure S2A and B**. A similar rise of RBF was observed in every rat studied and a similar increase in the magnitude of the diurnal rhythm of the RBF was observed in all rats. Shown in **Figure S3**, total urinary Na^+^ excretion increased 10-fold (from 0.9 ± 0.1 to 10.3 ± 0.4 μmol/min) consistent with the 10-fold increase in the percent of NaCl in the diet when switch from 0.4% salt to 4.0% NaCl.

**Figure 1.**
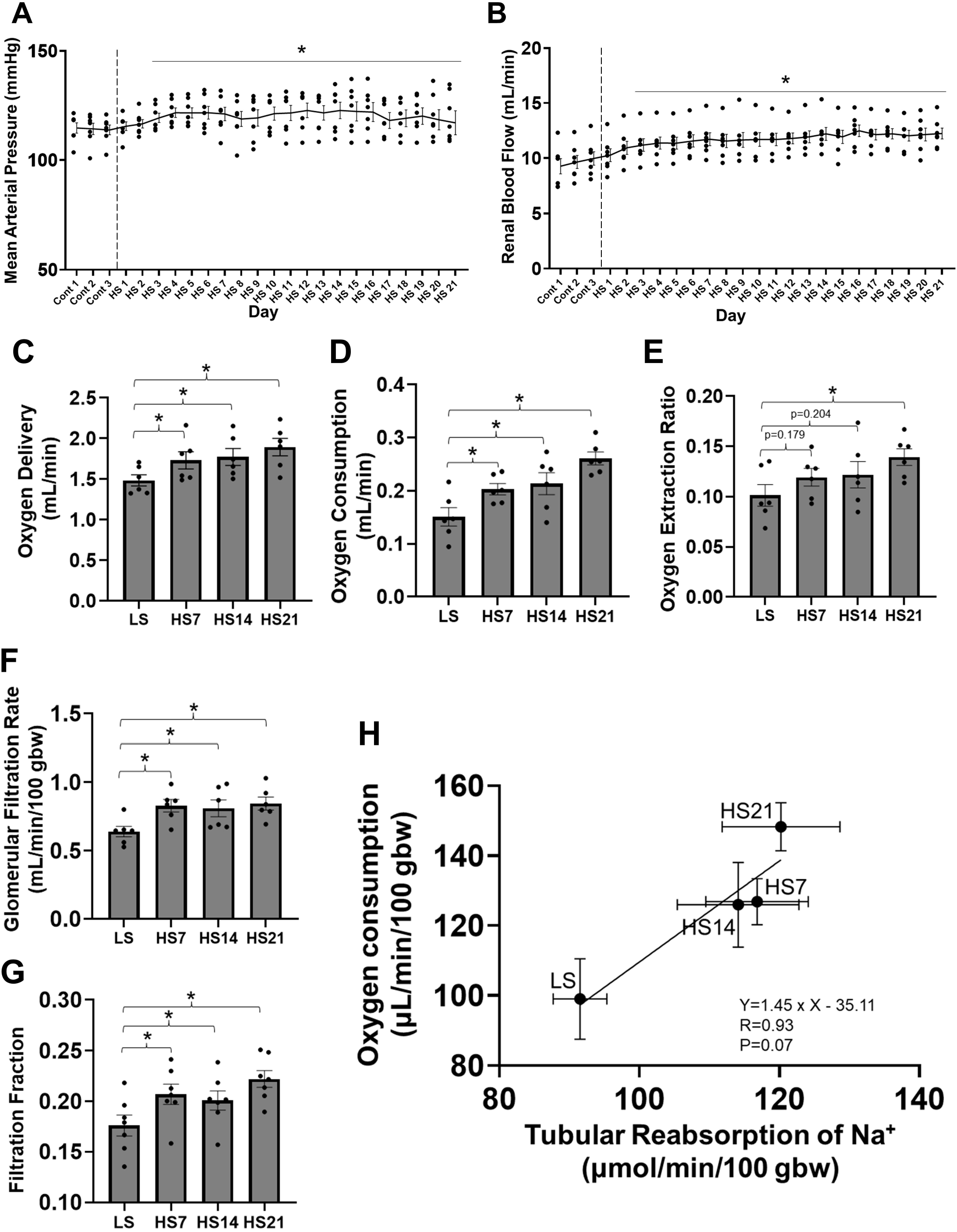
Physiological responses to the 4.0% NaCl diet. (A) Twenty-four hour/day average mean arterial pressure (MAP), (B) renal blood flow (RBF), (C) oxygen delivery, (D) oxygen consumption, and (E) oxygen extraction ratio obtained from same group of SD rats (n=7). (F) Glomerular filtration rate (GFR) measured in a separate group of rats (n=6), (G) Filtration fraction. (H) Correlation between O_2_ consumption and tubular reabsorption of Na^+^ determine using data from the latter two groups of rats. RBF and O_2_ data for single kidney were doubled for (G) and (H) to represent whole body values. One-way repeated measures (RM) ANOVA, Holm-Sidak for (A) – (G) and Pearson’s r correlation coefficient (H) were performed. Mean ± SEM and individual data. *p<0.05 vs Cont 3 (A), (B) or LS (C) – (G).

With the increase of RBF, the O_2_ delivery increased proportionally to RBF due to the unchanged O_2_ content in artery (**Table S4**) and remained elevated throughout the 21 days of the HS diet (**Figure 1C**). Both the calculated O_2_ consumption (**Figure 1D**) and extraction ratio (**Figure 1E**) rose progressively throughout the 21 days of the HS diet with the ratio increasing from 0.10 ± 0.01 to 0.14 ± 0.01 (p<0.05) by HS21. This trend was maintained when normalized by body weight (O_2_ delivery: p=0.186; O_2_ consumption and O_2_ extraction; p<0.05) (**Table S3**).

### Effects of high salt diet on GFR and calculated filtration fraction

In a separate group of rats, the total body GFR was determined in unanesthetized rats which underwent the same surgery (sham) as those with implanted renal flow probes and subjected to the same dietary protocol. As summarized in **Table S2** and **Figure 1F**, GFR increased from 0.64 ± 0.04 at LS to 0.83 ± 0.05 at HS7 (mL/min/100 g body weight) (p<0.05) and was thereafter maintained at that level throughout the study. The filtration fraction (**Figure 1G**), calculated using the RBF determined from the continuously recorded rat group, was increased from 0.18 ± 0.01 at LS to 0.22 ± 0.02 at HS7 and remained elevated at this level throughout the study. As determined from these data, a strong positive correlation was found between total tubular reabsorption of Na^+^ and O_2_ consumption as presented in **Figure 1H**.

### Effects of high salt diet on untargeted metabolomic profiles of cortical and outer medullary tissue

**Figure S4A** summarizes the total number of compounds detected in the renal cortical tissue (Cx) (5959) and in the outer medullary tissue (OM) (5785) combining the results detected in all modes. As illustrated, of the 5959 compounds in the Cx and 5785 in the OM, 2851 and 2620 are named compounds by Thermo Compound Discoverer (**Figure S4B**). Of all named compounds, 968 in Cx and 861 in OM were found in the Metaboanalyst 5.0 database (**Figure S4B**). Comparing LS values to those obtained after 14 days of the HS diet (HS14), it was found that 284 metabolites were significantly changed (raw p<0.05) in the Cx and an equal number (284) significantly changed in the OM. Comparing LS values to those obtained after 21 days of the HS diet (HS21), 438 were significantly changed in the Cx and 349 in the OM (raw p<0.05) (**Figure S4B**). P-value adjustments (Benjamini-Hockberg) reduced the chance of making Type-I errors and it was found that 138 named metabolites were significantly changed at HS14 compared to LS (adjusted p<0.05) and 229 in the OM. At HS21, 218 were significantly changed in the Cx and 225 in the OM.

A sparse partial least squares discriminant analysis (sPLS-DA) was carried out for these named compounds to visualize and assess similarities and differences of metabolites in response to the HS diet. For each of the 4 modes (C18 +/- and HILIC +/-), 5 components with 20 variables per rat were analyzed. As shown in **Figure 2A**, the analysis revealed that separate metabolic states were distinguishable within the Cx and OM tissues in response to the HS diet at both days 14 and 21. A clear distinction is observed between LS and HS14 and HS21 days of feeding.

**Figure 2.**
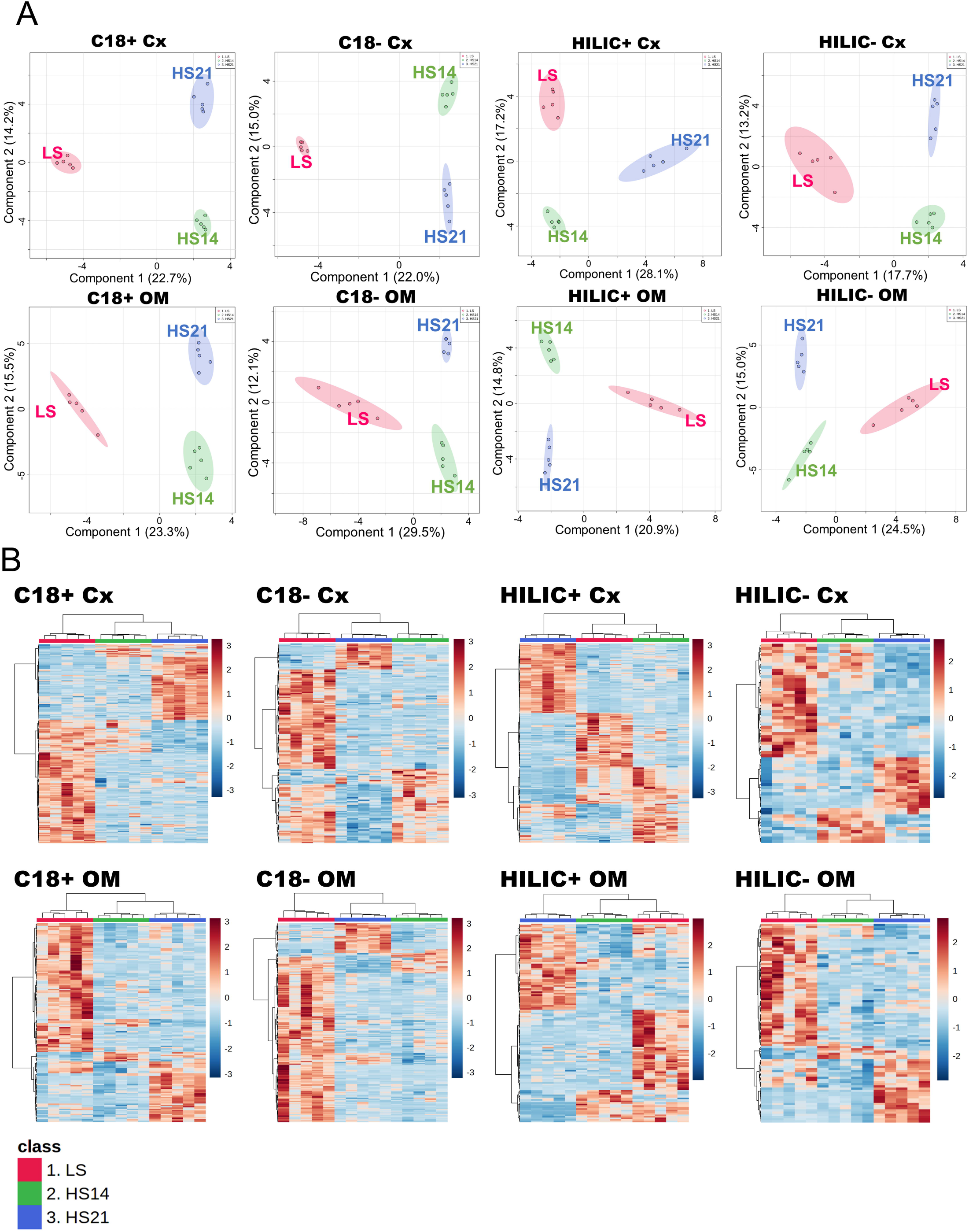
Sparse Partial Least-Squares Discriminant Analysis (sPLS-DA) of cortical (Cx) and outer medullary (OM) tissues in each of 4 modes. (A) Parameters for sPLS-DA are fixed to 5 variables per component and to 20 for validation yielding 5-fold difference in coefficient of variation. Ellipses represent 95% confidence region of a bivariate normal distribution. (B) For those compounds significantly affected by high-salt (HS) (ANOVA Fisher’s LSD p<0.05) a hierarchical cluster analysis was performed with a Euclidean distance measure and by the Ward algorithm and represented by heatmaps. Red: Low salt (LS), Green: 14 days of HS, Blue: 21 days of HS.

The significantly altered metabolites (ANOVA Fisher’s LSD p<0.05) determined in the C18+ mode were 321 of 1114 in Cx and 151/668 in OM. The significantly altered metabolites in the C18-mode were Cx 161/649 and OM 225/804; in the HILIC+ mode were Cx 241/565 and OM 129/805; and in the HILIC-mode were Cx 63/564 and OM 91/470. The heat maps of those metabolites are shown in **Figure 2B** in which the distinctive patterns observed by sPLS-DA were validated in the hierarchical clustering which was performed with a Euclidean distance measure and by the Ward algorithm.

A metabolite enrichment analysis was performed for those metabolites that were significantly changed (p<0.05) to prioritize and place them into known biological pathways as described by the small molecule pathway database (SMPDB). As shown in **Figure S5**, the “arachidonic acid metabolism” was significantly enriched in the Cx at HS14. In the OM, the “tyrosine metabolism”, the “lysine degradation”, and the “beta alanine metabolism” were significantly enriched at HS14. Notably, at HS21, no enrichment of any of the metabolomic pathways was found either in the Cx or in the OM.

### Effects of high salt upon cortical and outer medullary mRNA expression (mRNAseq analysis)

mRNAseq analysis was performed on the same tissue analyzed for metabolites as described above to validate identification of enriched metabolomic pathways using a denser genomic scale dataset (e.g., ∼3000 named metabolites vs. ∼30,000 gene transcripts). There were 32,545 genes identified by mRNAseq. Within that, adjusted p<0.05 by Benjamini and Hochberg’s test, comparing LS to HS14, 497 significantly increased and 422 decreased in Cx. Comparing LS to HS21, 3044 increased and 2917 decreased in Cx., In the OM, comparing LS to HS14, 91 increased and 22 decreased and comparing LS to HS21, 555 increased and 165 decreased.

The pathway analysis of mRNAseq data (KEGG) **(Figure 3)** found that genes most upregulated in the Cx tissue of SD rats fed HS diet were those related to the “Signaling molecules and interaction pathway”, the “Immune system pathway” (red bars), and related pathways including “Cytokines receptors”, “Chemokine signaling”, “NF-kappa B signaling”, “Th17 cell differentiation”, “T cell signaling”, etc. Those pathways most downregulated were those related to metabolism (blue bars) including “TCA cycle”, “Fatty acid degradation”, “Valine, Leucine and Isoleucine degradation”, “Glycine, Serine and Threonine metabolism”, “Carbon metabolism”, etc. In the OM tissue, the upregulated pathways were also largely related to the immune system, but interestingly fewer pathways of metabolism were found to be downregulated with the HS diet.

**Figure 3.**
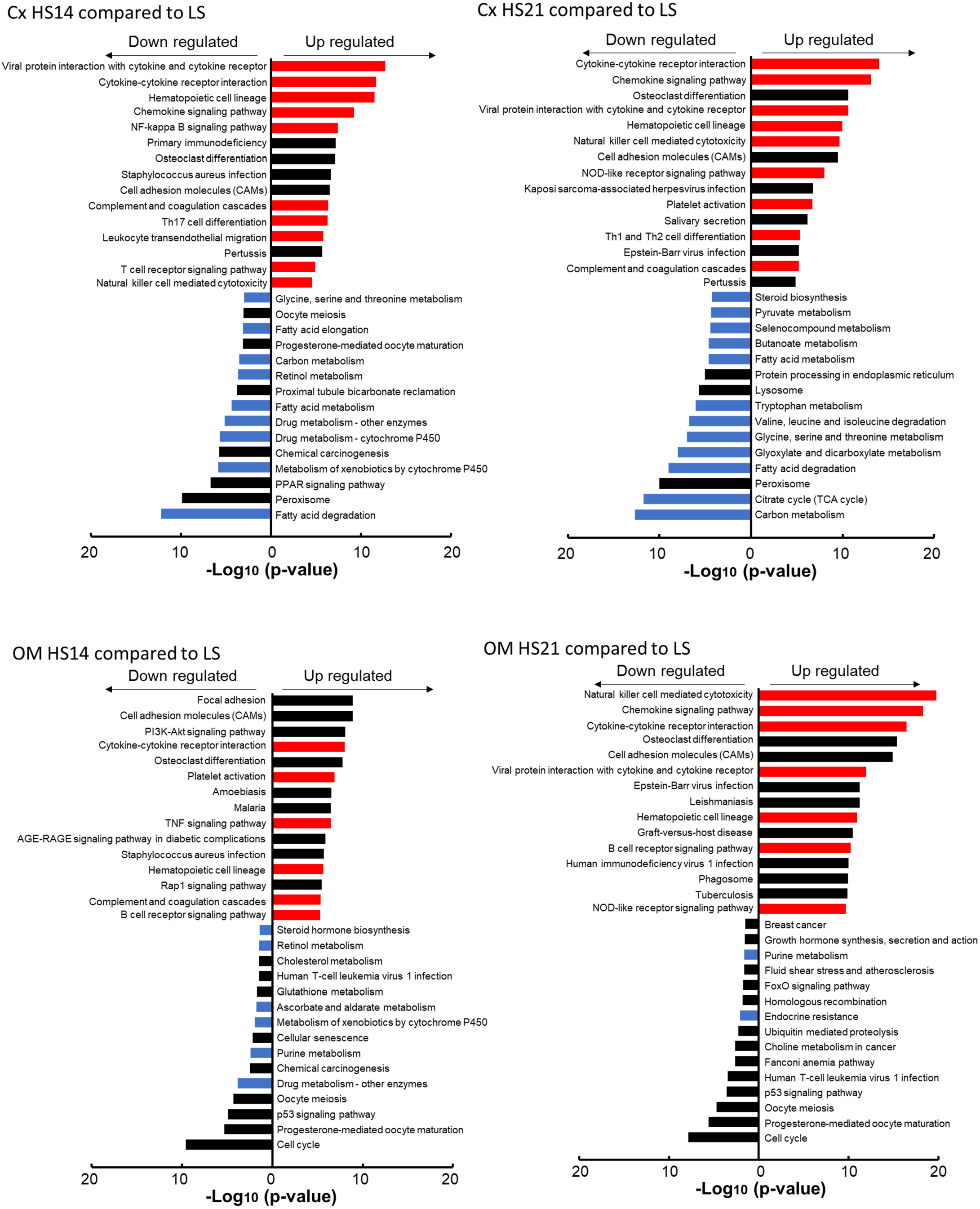
Results of the pathway analysis of the mRNAseq data showing the top 15 KEGG pathways (in order of -log_10_ p-values) that were either significantly increased (“up regulated”) or significantly decreased (“down regulated”) in cortex (Cx) and outer medulla (OM) at either HS 14 or HS 21 compared to gene expression with the LS diet. Red bars denote “Signaling molecules and interaction pathway” and “Immune system pathway (KEGG map ID 04XXX)”. Blue bars denote “Metabolism pathway (KEGG map ID 00XXX, 01XXX)” (August 2022).

After excluding genes with a count of less than 100 in the mRNAseq data, the top 20 and bottom 20 with the largest fold change (FC) in HS compared to LS are shown in **Table S5**. In addition to some genes showing immune system upregulation (i.e. *C1s, C3, RT1-Db1, Itgal*, etc), genes with large FCs in Cx were also found in the *Mthfr* gene whose polymorphisms are related to hypertension^34^, *Nrep* also known as *P311* stimulates translation of TGFβ and is related to tissue fibrosis^35^, and *Slc16a1* also known as *Mct2* is a proton-linked monocarboxylate transporter^36^. Immune system related genes were also upregulated in OM.

### mRNA expression of cortical tubular transporters

Of special mechanistic interest were the changes in gene expression of tubular transporters affected by the HS diet. **Figure S6** shows the Cx tissue mRNA expression of genes encoding transporters for glucose and amino acids that tended to be downregulated at HS14 with most of these reaching statistical significance by HS21. This includes amino acid transporters (*Slc1, Slc3, Slc7*), sodium-glucose transporters (SGLT isoforms *Slc5a1, Slc5a2*), urate (*Slc22a12*) and lactate transporters (*Slc5a12, Slc5a8*). It was found that the glucose transporter 2 (GLUT isoforms *Slc2a2*) were down regulated at HS21 while expression of *Slc2a1* was increased. Although it is unclear which of these would predominate functionally, it is evident that the upregulation of the GLUT transporters would be consistent with a greater release of glucose and glycolysis products being released into the interstitial space. Also of note, monocarboxylate transporters (MCT) and Na^+^-K^+^-ATPases were upregulated by the HS diet which is consistent with enhanced proximal tubule Na^+^ reabsorption necessitating greater utilization of ATP for active transport. Sodium transporters and channels including NHE3 (*Slc9a3*), NKCC2 (*Slc12a1*), NCC (*Slc12a3*) and ENaC (*Scnn1*) were upregulated. On the other hand, the effect of HS on the OM transporter genes was modest (**Figure S7)**.

In addition, proteolysis related genes are picked up in **Figure S6** as well. There are myriads of proteolytic enzymes expressed in the kidney^37, 38^, and those are effected by HS diet in Cx tissue. Genes encoding protease-activated receptors (*F2r*)^39^ are upregulated by HS, whereas megalin (*Lrp2*) and clathrin (*Cltc*) were reduced. Interestingly, within cathepsin encoding genes, *Ctsa* and *Ctsb* which mainly expressed at proximal tubule (PT) S1 were downregulated and *Ctsc* and *Ctsd* which mainly expressed at distal tubule (DT) and connecting tubule (CNT) were upregulated.

### Omic integration with statistical mapping and validation of metabolomic and mRNAseq data

Data obtained from the metabolomics and mRNAseq analysis can be effectively utilized in several ways. First, to validate against each other the predicted pathways that appear to be most affecting metabolic functions when fed a high salt diet. Second, to identify pathways that were not of obvious importance from the metabolomic analysis given the limited number of compounds that can currently be identified by a global mass spec analysis.

Figure 4. summarizes the integrated metabolomic and mRNA expression data related to mitochondrial energy production in the Cx sample. By HS21, both metabolites and genes encoding the major enzymes of the TCA cycle were found to be generally downregulated including reductions of citric acid, succinic acid, and fumarate. While mRNAs encoding enzymes in the TCA cycle were downregulated overall, some of mRNAs encoding proteins that make up the electron-transfer complex were not (**Figure 4**). Especially, many genes encoding for many proteins of complex V including *Mt-atp8* were upregulated at HS21 while the expression of others was reduced. It was also found that the gene encoding uncoupling protein 2 (*Ucp2*) was upregulated by HS.

Several other important pathways were enriched in the metabolomic enrichment analysis (**Figure S8**). Specifically, the “Arachidonic acid metabolism” pathway in the Cx (**Figure S8A**) and the “Lysine degradation” pathway in the OM (**Figure S8B**) were altered by the HS diet. As shown in **Figure S8A**, the metabolomic analysis in the Cx found that arachidonic acid was significantly increased at HS14, together with downstream metabolites 8-HETE and thromboxane B2 (TXB2). The general upregulation of this pathway is reinforced by enhanced expression of most of the genes on arachidonic acid metabolism pathway which were upregulated on HS14 and HS21 compared to expression observed with the LS diet. The notable exception to this were the *Cyp-4* genes important in the production of 20-HETE which has both pro- and anti-hypertensive actions resulting from modulation of vascular and tubular functions of the kidney^40^. **Figure S8B** illustrates the “Lysine degradation” pathway in which many key elements were found in the metabolomic analysis to be downregulated in the OM as found at days HS14 and HS21. This included reduction of tissue lysine itself and reduced levels of α-ketoglutarate, glutamate, and allysine. However, since lysine was significantly reduced, a reduction in the metabolites derived from lysine would be expected to be also reduced since no significant changes in the enzymes in this pathway were observed by the mRNAseq analysis.

**Figure 4.**
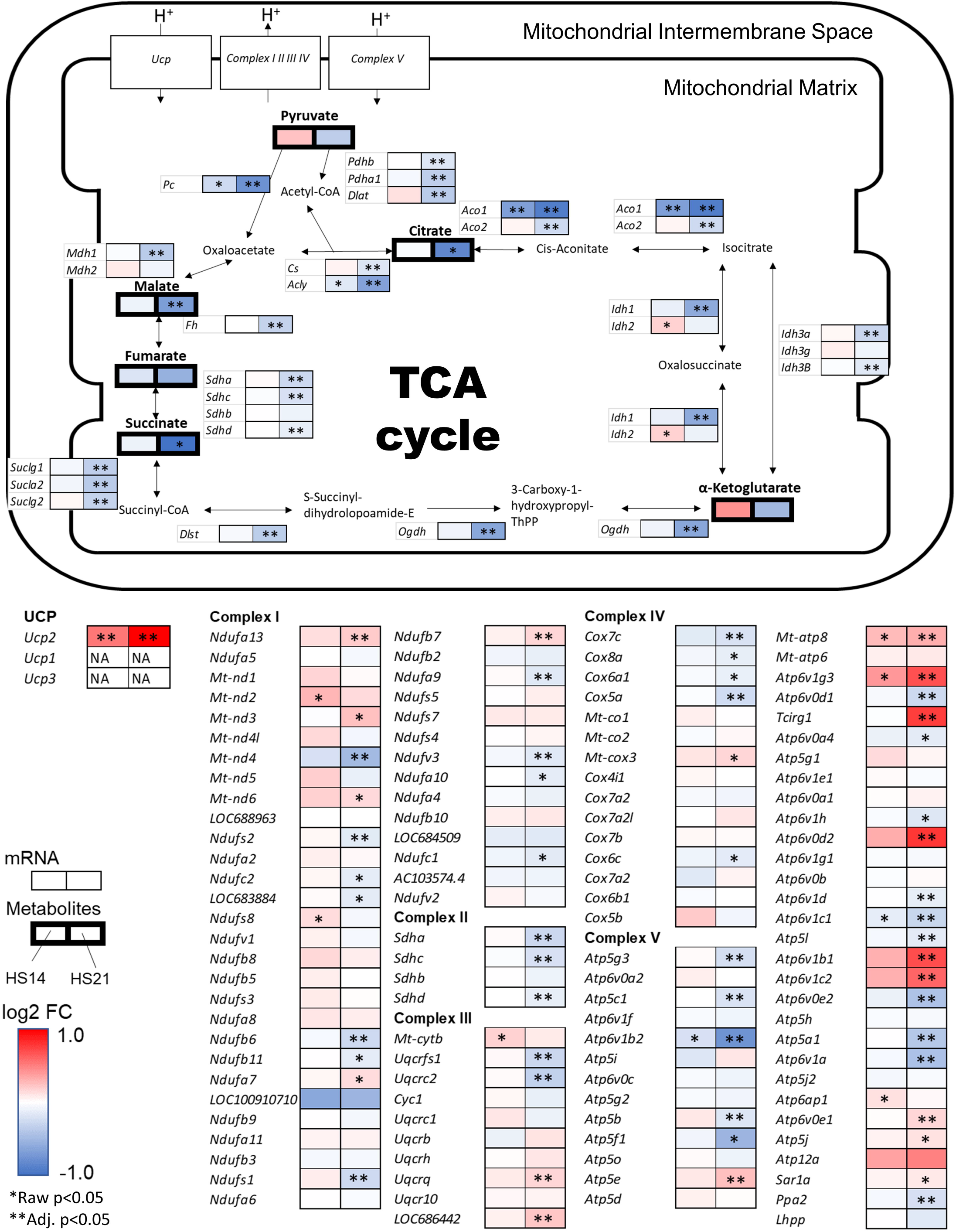
Integrated figure of TCA cycle in cortex Log_2_ fold change (FC) of high salt (HS) to low salt (LS) are represented in color. The left boxes are FC of HS14 to LS and the right boxes are FC of HS21 to LS. Thin boxes represent mRNA and thick boxes represent metabolites. Red denotes increase in expression and blue denotes decrease in expression. *Raw p<0.05 in t-test for metabolomics and in DESeq2 for mRNAseq, **Adj. p<0.05 in Benjamini and Hochberg. (KEGG map ID 00020, 00190 last update: July/28/2022)

### Effects of the high salt diet upon glycolysis in the renal cortex

As illustrated in **Figure 5A**, the Cx levels of glucose tended to be reduced by day 14 of the HS diet and were significantly reduced by day 21 of the HS diet. A seemingly compensatory response is reflected by the observation that many genes encoding glycolytic enzymes were upregulated at HS14 and even more so at HS21. Significant increases (p<0.05) were found in hexokinase isoforms (*Hk1, Hk2, Hk3*) required for conversion of glucose to glucose 6-P, and in phosphofructokinase isoforms (*Pfkl, Pfkp, Pfkm*), required to convert glucose 6-P to fructose 1,6-BP. A reduction of glyceraldehyde 3-P and 3-phosphoglycerate was found at HS21 and several isoforms of aldo-keto reductase (*Aldoa, Aldob, Aldoc*) and glyeraldehyde-3-phosphate dehydrogenase (*Gapdh*) were found to be reduced at HS21. Increases were also found in pyruvate kinase (*Pkm, Pklr*). The Cx pyruvate levels tended to be increased at HS14 but did not differ from levels observed with LS at HS21. Lactate dehydrogenases (*Ldha, Ldhb, Ldhc*) were also upregulated, but the tissue lactate did not differ from levels observed with LS at either HS14 or HS21.

**Figure 5.**
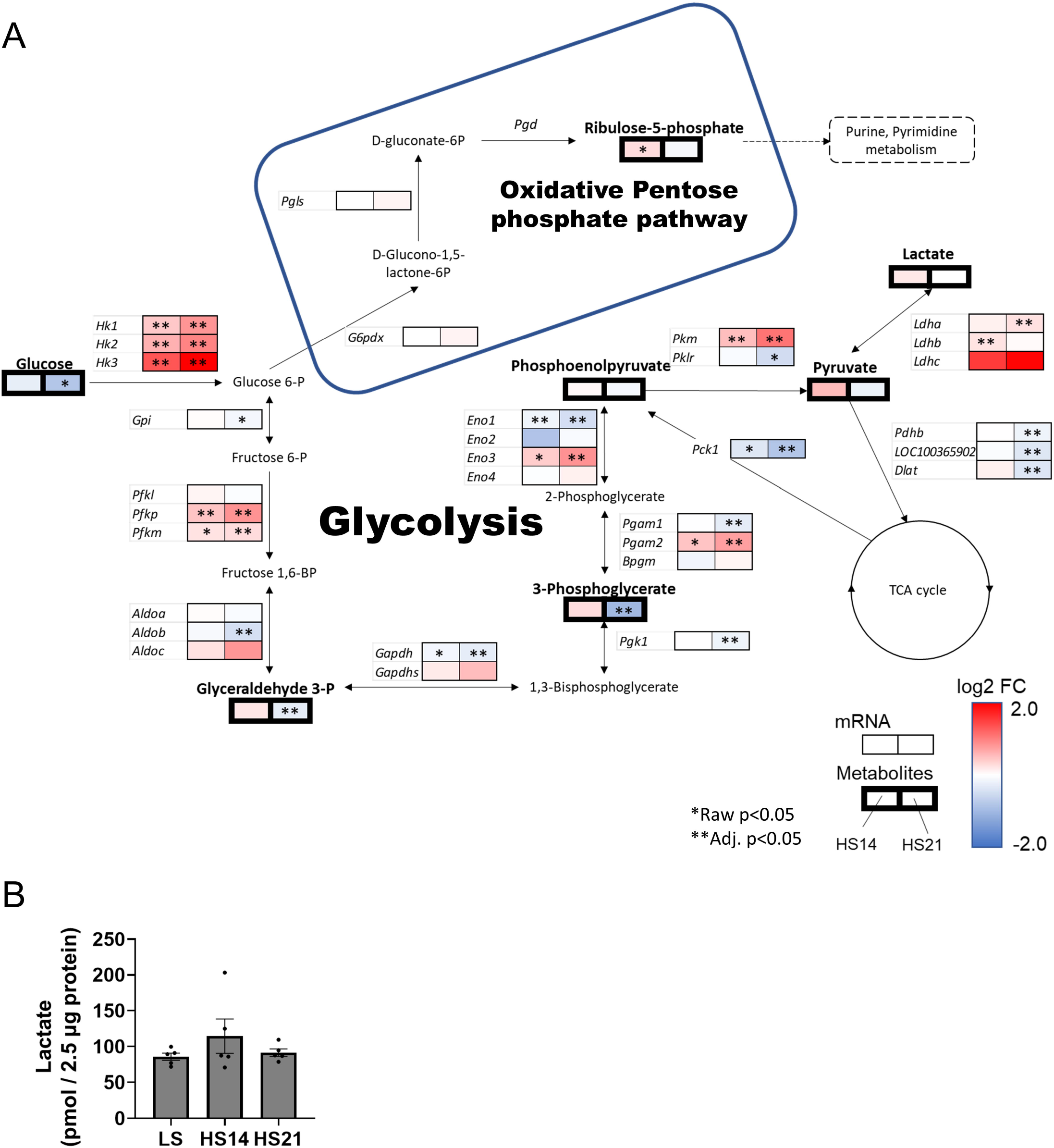
Integrated figure of glycolysis in cortex (A) Log_2_ fold change (FC) of high salt (HS) to low salt (LS) are represented in color. The left boxes are FC of HS14 to LS and the right boxes are FC of HS21 to LS. Thin boxes represent mRNA and thick boxes represent metabolites. Red denotes increase in expression and blue denotes decrease in expression. The frame denotes oxidative pentose phosphate pathway. *Raw p<0.05 in t-test for metabolomics and in DESeq2 for mRNAseq, **Adj. p<0.05 in Benjamini and Hochberg. (KEGG map ID 00010 last update: May/7/2020, 00030 last update: December/13/2022). (B) Validation of lactate concentration in cortical tissue by fluorometric assay kit. Mean ± SEM and individual data. One-way ANOVA, No significant difference between groups.

The relationship of glycolysis with the oxidative pentose phosphate pathway (PPP) is also illustrated in **Figure 5A**. A moderate increase (p<0.05) of ribose-5-phosphate was found at HS14 indicating a greater oxidation of glucose via this pathway which at this time would yield greater NADPH to scavenge ROS. However, at HS21 ribose-5-phosphate was no longer found to be elevated. As activation of glycolysis was of particular interest, lactate concentration in the Cx tissue was validated by fluorescent lactate analysis which showed the similar trend as metabolomics data (**Figure 5B**). Further arteriovenous flux analysis is discussed below.

In general, it appears that metabolites of the glycolytic pathway in the Cx may be initially elevated at HS14 but by HS21 appear to be reduced to levels similar to or less than that observed when fed the LS diet. As noted in **Figure S6**, gene expression of the PT apical membrane SGLT transporter *Scl5a1* isoform was increased at HS14 (p<0.05) but not at HS21. A significant reduction of the *Scl5a2* isoform was found at HS21 suggesting there could be a reduced luminal uptake of glucose in the PTs, perhaps contributing to the reduced Cx glucose levels. Gluconeogenesis may be expected to be suppressed since phosphoenolpyruvate carboxykinase 1 (*Pck1*) that acts as the rate limiting enzyme in gluconeogenesis was reduced (**Figure 5A**).

Many genes involved in the glycolytic and TCA cycle were altered in Cx, while less significant changes were observed in OM (**Figure S9**).

**Figure S10** summarizes the genes and metabolites related to the malate-aspartate shuttle. The mRNAseq analysis confirmed that this pathway exists in the Cx, but since gene expression was not assessed separately in the cytoplasm and mitochondria one cannot determine whether the expression of enzymes that directly supply protons, such as *Mdh*, were altered. Malate levels were reduced at HS21 (p<0.05) as was *Slc25a11* gene expression which codes for the mitochondrial carrier that transports malate across the IMM. It is interesting that *Slc25a12* which codes for the protein that transports aspartate across the IMM to the intermembrane space was significantly increased at HS21, but the relevance of this is unclear.

**Figure S11** summarizes the genes and metabolites related to urea cycle and nitric oxide production in the renal Cx. It was found in the Cx that citrulline and arginine were significantly reduced together with a reduction in *Nos1* mRNA expression at HS14. Aspartate was found to be elevated perhaps representing a compensatory response. By HS21, arginine had returned to levels observed with LS and was associated with elevations of *Nos3* which appears to represent a compensatory response. *Arg2* mRNA expression was also elevated at HS21 which could also drive an increase of urea production although urea was not measured by the mass spec analysis.

### Arterial and venous metabolites and flux determinations

As shown in **Figure S12**, a total of 3137 of compounds in plasma were detected when all modes were combined. Of these,1531 are named compounds.

We first analyzed the metabolomic profiles in the artery (Art) and renal vein (RV) separately. It was found that the Art (**Figure 6A**) and RV metabolomes (**Figure 6B**) as analyzed by a sparse partial least squares discriminant analysis (sPLS-DA) exhibited a clear shift in the metabolic states at HS7 and HS14. It is also evident that by HS21, the metabolic state was more similar to that found when rats were fed the LS diet representing a return to “normal” although clear differences are seen in component 2 metabolites in both Art and RV samples. This return of the metabolomic profiles toward that of the LS state after 21 days of the HS diet is also clearly seen in the associated heat maps representing those metabolites that were significantly changed by the HS diet for each of the designated modes (ANOVA Fisher’s LSD p<0.05). Specifically, the significantly altered metabolites in different modes are as follows: C18+ Art 96/808, C18+ RV 65/808, C18- Art 35/454, C18- RV 38/454, HILIC+ Art 31/387, HILIC+ RV 40/387, HILIC- Art 19/122, HILIC- RV 15/122.

**Figure 6.**
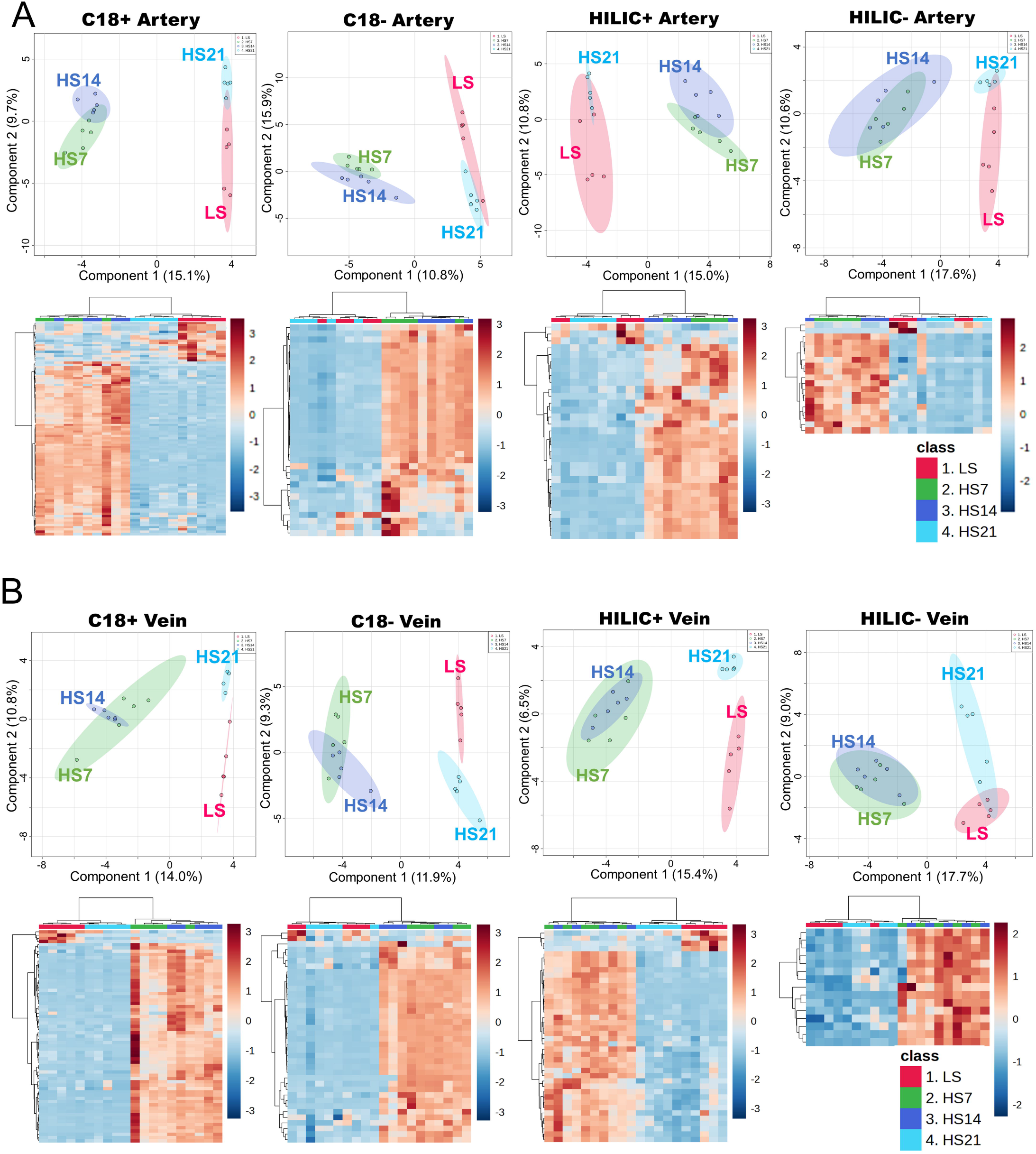
sPLS-DA and heatmap of tissue Sparse Partial Least-Squares Discriminant Analysis (sPLS-DA) of arterial and venous plasma in each of 4 modes (A). Parameters for sPLS-DA are fixed to number of components: 5, variables per component: 20 and validation method: 5-fold CV. Ellipses represent 95% confidence region of a bivariate normal distribution. Compounds which are significantly affected by high-salt diet (HS) (ANOVA Fisher’s LSD p<0.05) are performed hierarchical cluster analysis with a Euclidean distance measure and by the Ward algorithm and represented by heatmaps (B). Red: Low salt (LS), Green: 7 days of HS, Blue: 14 days of HS, Light blue: 21 days of HS.

Focusing the analysis on the differences between the Art and RV of those named compounds in response to the HS diet it is seen in **Figure S13A** that by sPLS-DA analysis that separations were found between Art-Rv metabolites in the C18+ and HILIC+ modes comparing LS and HS7, HS14 and HS21. However, there was no clear separation found between the days of the HS diet. As illustrated in **Figure S12B**, within the named 1531 compounds, 546 compounds were contained in the Metaboanalyst 5.0 database. Of these, 132 compounds (**Table S6**) were significantly (raw p<0.05) changed over time by HS as determined using linear models with covariate adjustments^41^ on Metaboanalyst 5.0. Enrichment analysis of those metabolites revealed the significant enrichment of “oxidation of branched chain fatty acids” and “carnitine synthesis” (**Figure S13B**) including the metabolites such as carnitine, acetylcarnitine, lysine and α-ketoglutarate.

The urine metabolomic features are summarized in **Figure S14** showing that a total of 10,241 compounds were identified in the urine sample from the four modes (C18+/- and HILIC+/-). Of these, 4980 were named compounds and after removing those compounds not found in plasma and duplications, there remained 749 compounds. Of these, only 367 were found to match the Metaboanalyst 5.0 database which were used for the final flux analysis. Those metabolites in the urine for which the urine excretion exceeded the total kidney filtration fraction are shown in **Table S7**. With LS feeding, 19 metabolites were found to be excreted in excess of the filtration fraction. Interestingly, the number of metabolites excreted more than filtration fraction kept increasing over time (29 at HS7, 37 at HS14, and to 42 at HS21) and began to include uremic toxins such as creatinine, indoxyl sulfate, and hippuric acid.

Calculated metabolite fluxes comparing those determined in rats fed LS to those at HS7, HS14 and HS21 are shown in **Figure 7**. In these individual graphs, metabolite fluxes are represented from two perspectives. First, whether the metabolites were released or taken up. For this purpose, the 95% confidence interval of the mean was calculated and if “0” was not contained within this confidence interval it indicates whether a compound is either released or taken up by kidney (p<0.05). Second, the graphs reflect whether the fluxes changes over time, for which a one-way repeated measures ANOVA post hoc Holm-Sidak was performed comparing all times to the LS fed state (p<0.05 indicated by *). The analysis indicates that the flux of some of the important carbohydrates utilized for both glycolysis and the TCA cycle were modified by HS intake (see graphs in blue boxes). Specifically, it is seen that in the LS state, glucose was being generated in the kidney (p<0.05) which was not observed during the days of HS feeding. Lactate release became significant (p<0.05) at days 14 and 21 of the HS diet with a similar trend in pyruvate. Plasma lactate concentration was validated by fluorescent assay kit and similar tendency was observed (**Table S8**). There was a greater uptake of the important TCA cycle intermediate α -ketoglutarate (p<0.05) in the LS fed rats which was no longer apparent with HS feeding. An uptake of citrate was found with the LS diet and this positive uptake was sustained throughout most of the periods of HS feeding (p<0.05) except at HS14. Gluconeogenic amino acids (see graphs in orange boxes) including asparagine, tryptophan, phenylalanine, valine, arginine, glutamate, histidine, and proline were released in significantly greater amounts (p<0.05) during various days of the HS diet. The ketogenic amino acid lysine was also released from the kidney at HS14 (see graph in green box).

**Figure 7.**
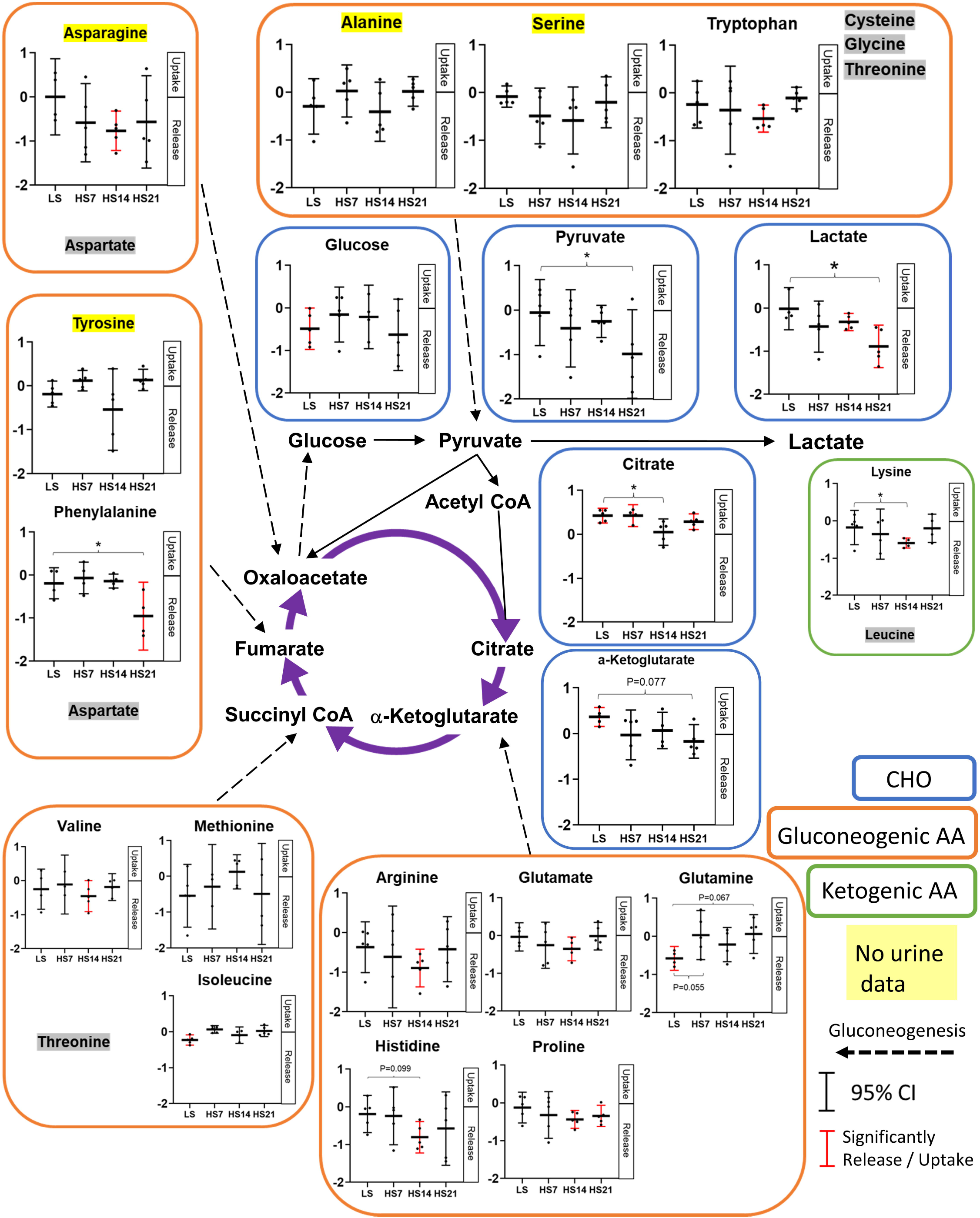
Calculated metabolic fluxes Mean ± 95% confidence interval of mean and individual data are shown in each graph. Horizontal line is date, vertical line is flux (calculation written in methods. no unit). Red graph denotes the significantly taken up or released in each time point. *p<0.05 vs low salt (LS), one-way RM ANOVA, Holm-Sidak. Blue boxes: carbohydrates (CHO), orange boxes: gluconeogenic amino acid (AA), green boxes: ketogenic AA. Purple arrows denote TCA cycle and dash arrows denote gluconeogenesis. Highlighted by yellow are metabolites which are not detected in urine (i.e. only arterial and venous plasma are used for flux calculation). Highlighted by gray are not detected in plasma. (Adapted the figure of Stryer Biochemistry. 7th edition (2012) ^99^)

## Discussion

The kidneys play a crucial role in eliminating excess salt in the diet and maintaining homeostasis in the body. In salt sensitive individuals with reduced sodium excretory function BP increases when fed a high salt (HS) diet which in turn leads to vascular, cardiac and kidney dysfunction and injury. Although excess salt intake is less likely to produce hypertension in individuals with low salt sensitivity^42-44^, the present study finds that a HS diet does have a significant effect on kidney metabolism even in normal Sprague Dawley (SD) rats in which minimal hypertension is observed when fed a HS diet.

### Effect of excess salt on renal hemodynamics and O_2_ utilization in SD rats

As determined by continuous 24 hr/day monitoring, the SD rats as expected showed only a slight increase in the average daily MAP (∼5 mmHg) in response to the HS diet. By comparison, a relatively much larger increase in RBF was observed. An even greater increase of GFR was observed consistent with the previous observations^45^ although the mechanism for this are unclear. Nevertheless, this resulted in the significant increase in the calculated filtration fraction (FF). Although high GFR salt sensitivity is thought to be associated with greater susceptibility to progression of renal dysfunction^46-48^, it is clear that SD rats possess compensatory mechanisms that enable them to compensate and prevent the injurious effects of a HS diet. This is in stark contrast to Dahl SS rats, which were generated by selective breeding of SD rats and whose GFR is reduced by the second week of HS feeding^27^. Increased filtration fractions with HS intake have also been observed in salt sensitive humans (increase in MAP by 8 mmHg, increase in FF by 0.04) and women using oral contraceptives (increase in MAP by 1-2 mmHg, increase in FF by 0.02) ^49, 50^. As O_2_ content in arterial blood did not change over time with the HS diet, O_2_ delivery increased in proportion to RBF. On the other hand, O_2_ consumption increased proportionally greater than the increase in the delivery. Although a HS diet was reported to decrease the tubular O_2_ consumption in isolated micro-dissected renal tubules from mice^51^, this does not appear to reflect in vivo responses where tubular O_2_ consumption is altered by many factors including GFR and RBF that must be taken into account^52^. Although the correlation between Na^+^ reabsorption and O_2_ consumption in the kidney is well recognized as determined under a variety of conditions^53 30^, to the best of our knowledge, this is the first report that has evaluated this relationship in the unanesthetized freely moving animal repeatedly. In previous In vivo experiments with dogs, renal Na+ reabsorption was reported to be 20 mol per mol of O_2_ ^12, 53^. Our data at LS is 20.7 Na/O_2_ (assuming 22.4 L/mol of O_2_), which is close to the previously reported value. It is noteworthy that the value of Na/O_2_ did not significantly change even under the HS (18.2 at HS21). Given that 99% of the filtered Na^+^ is reabsorbed, the tubular load is primarily dictated by increase in GFR so an increase of GFR would be expected to increase the tubular workload and O_2_ consumption.

Salt administration altered the expression of many genes encoding cortical tubular Na^+^ transporters which was especially evident at HS21. The enhanced expression of Na^+^ transporters in the cortex may have affected metabolic changes. It was found that Na^+^ transporters and channels were generally upregulated whereas sugar and amino acids transporters were found to be downregulated. As illustrated in **Figure S6**, in cortical tissue (Cx), increased gene expression was found of NKCC2 (*Slc12a1*) which is expressed in cTAL, NCC (*Slc12a3*) which is expressed in the aldosterone sensitive distal collecting tubules (DCT), ENaCa (*Scnn1a*) which is expressed in the DCT and cortical collecting ducts (CCD), and NHE3 (*Slc9a3*) which is expressed in proximal tubules (PT)^54^. Together, the increased mRNA expressions of the transporters would be expected to increase both proximal and distal tubular Na^+^ reabsorption^55, 56^. These observations are consistent with the conclusions reached by Udwan et al.^51^ that the fractional reabsorption of Na^+^ is distributed differently along the tubule as determined by dietary Na^+^ intake.

### Change in metabolites and gene expression by the high salt diet

Despite the current progress in mass spectrometry and the ability to detect more than 5,000 metabolites within biological samples, the number of annotated compounds reduces that number to several thousand and of those the assignment to known biochemical pathways is a limiting factor when compared to those obtained from mRNAseq analysis in which nearly 30,000 known protein-coding genes can be mapped to less than 1,000 biochemical metabolites represented in the KEGG pathway maps. In the present study we have utilized the combined strength of large-scale transcriptome sequencing (mRNAseq) with global profiling of metabolites in which the integrated analysis has identified many pathways of metabolism in which statistically significant differences to salt diet were obtained. Even those which did not reach statistically significant differences for the metabolomic analysis were of great utility when changes were consistent with pathways found of importance in the mRNAseq analysis.

### Compensatory mechanisms to protect from hypertension and kidney injury in SD rats

One of the important underlying questions is what are the underlying compensatory mechanisms that protect the SD rat from hypertension and kidney injury when fed a HS diet. Although the specific answer to this question is not provided by the present analysis, enormous changes are occurring in the kidney metabolism and in many of the molecular and biochemical pathways that affect major functions of the kidney and inflammatory pathways of tissue injury. Inflammatory pathways are stimulated by ROS which is generated during the process of oxidative phosphorylation. Several pathways work as ROS scavengers are found in this study. First, oxidative pentose phosphate pathway (PPP) might be upregulated which is evident from increase in metabolites, Ribulose-5-phosphate. PPP generate NADPH which is required for antioxidant system^57^. Second, *Ucp2* mRNA expression elevated by the HS diet, which is also a ROS scavenger^58^. Activation of *Nos3* mRNA expression was also found. NOS3 generate nitric oxide and work as a ROS scavenger^59^. These might interact with each other to scavenge ROS and protect kidneys from damage.

The results of this study show that the profile of metabolites both in the kidneys and in systemic (i.e. arterial plasma) changed over time in response to the HS diet as it was found that considerable changes occurred in both arterial and renal venous blood metabolic profiles. It is very interesting, however, that although significant changes were observed in the plasma metabolic profiles at HS days 7 and 14, the general metabolic profile returned in each of these rats to one similar to that observed with the LS diet by HS day21. This is in contrast to the profiles obtained from the Cx and OM tissue analysis which were markedly changed over the 21 days of the HS diet and did not return to LS levels. This raises the interesting question of whether extrarenal changes in metabolic function might play an important role in normally protecting the kidneys from the injurious effects of a HS diet and from organs these signals might arise. There are reports that a HS diet alters the gut microbiome^60, 61^ and liver metabolism^62^ which need to be explored in greater depth.

### Effects of the high salt diet on arachidonic acid metabolism pathway

Important effects of a HS diet on the arachidonic acid (AA) pathway were identified by both metabolomic and mRNAseq analysis which found the upregulation of many elements of this pathway significantly altered in the Cx at HS14 with a tendency to return toward LS levels at HS21 (**Figure S8**). It is well recognized that arachidonic acid is a major component of cell membrane phospholipids in the kidney which is metabolized by cyclooxygenase (COX), cytochrome P450 monooxygenase (CYP450), lipoxygenase (LOX) and leukotrienes (LTs) enzymes. COX production of prostaglandins (PG) and LTs leads to inflammatory injury in the kidney. CYP450 production of hydroxyeicosatetraenoic acids (19-HETE and 20-HETE) play important roles in tubular ion transport and in modulating tubuloglomerular feedback to regulate the load on the glomerulus^63^. It is interesting that despite increased expression of AA in the Cx, reduction of *Cyp4a1* and *Cyp4a2* genes was observed. This may reduce expression of 20-HETE which is known to reduce renal vasoconstriction and renal vascular responses to angiotensin II, endothelin, norepinephrine, nitric oxide and carbonmonoxide^64^ and play a key role in kidney damage during the inflammatory process. The effects of HS on the AA pathway have been found to contribute importantly to tubular transport, BP salt-sensitivity, and kidney injury in the Dahl SS rat model of hypertension^65-68^. We also observed a significant increase of TXB2 in the renal Cx at HS14 which appeared to be attenuated by HS21. TXB2 is an inactive metabolite of thromboxane A2 (TXA2) which is a potent vasoconstrictor and can lead to loss of renal structural integrity and inflammatory damage to the kidney^69, 70^.

### High salt downregulates the TCA cycle and upregulates glycolysis

There was a marked difference between the effects of the HS diet upon the metabolomic profiles of the Cx and OM. The Cx clearly showed major changes in the metabolic profiles while few changes were seen in the OM in response to the HS diet (**Figure 3**). One of the most interesting changes found in the Cx was related to the TCA cycle which at HS21 exhibited reductions in citrate, pyruvate, α-ketoglutarate, succinate, and in the mRNA expression of nearly all of the enzymes controlling the activity of the TCA cycle. Conversely, glycolysis appears to be upregulated indicated by related enzymes including hexokinases, pyruvate kinases, and lactate dehydrogenases.

Proximal tubules are thought to have limited capacity for glycolysis with energy needs being met by oxidative mitochondrial metabolism making them susceptible to damage with acute reductions of kidney perfusion^71^. It was surprising to find that the HS diet upregulated glycolytic gene expression in Cx in SD rat (**Figure 5**). However, downregulation of the TCA cycle and upregulation of glycolysis has also been observed in Dahl SS rats fed a HS diet^72, 73^. SD rats appear to possess counterregulatory pathways that protect the kidneys from the injurious effects of HS diets that occur in SS rats.

It is relevant that the HS diet did not produce major changes in the metabolomic profiles of the OM of SD rats. This strain to maintain normal levels of renal medullary blood perfusion. SS rats have been found to exhibit a rapid 30% reduction of medullary blood perfusion during the first week of a HS diet^27^ and we have found in a proteomic study of isolated mitochondria of these rats a down regulation which is not observed in salt-insensitive consomic SS.13^BN^ rats^74^. SS rats also exhibit a significant increase in total renal vascular resistance (RVR) when fed a HS diet while salt-insensitive rats (consomic SS.1^BN^ rats) in contrast to a rather reduction in RVR in salt-insensitive SS.1^BN^ and SD rats in the present study^75^. Protection from renal ischemia, especially in the renal OM of SD rats may preserve metabolic functions of the mTAL as suggested by an absence of a down regulation of the metabolism pathways when fed a HS diet (**Figure 3**). The downregulation of the TCA cycle proteins has been observed in mitochondria of isolated mTAL of SS rats^74^. Several studies from our laboratory have shown that reduction of medullary blood flow in the SD rat with chronic medullary infusion of H_2_O_2_ or an SOD inhibitor (DETC) result in a salt-sensitive form of hypertension^76, 77^. So too, reduction of renal medullary oxidative stress in SS rats by intrarenal infusion of L-arginine reduces salt induced hypertension in SS rats^78^. The lack of significant changes in metabolism-related genes in OM may reflect salt insensitivity in SD.

### Paradoxical relationship between kidney O_2_ consumption and TCA cycle activity

One of the most interesting observations of the present study was the seemingly paradoxical phenomenon of an increase in kidney O_2_ consumption and energy usage in face of a reduction in the TCA cycle activity. What is the source of this additional energy production and O_2_ usage? The data indicate that with the down regulation of the TCA cycle with the HS diet, glycolysis became a dominant source of energy production despite increased RBF increased O_2_ extraction. The increase of renal venous lactate (**Figure 7**) is consistent with increased activity of the glycolytic pathway. Aerobic glycolysis which was originally described in cancer cells by Warburg in 1921^79^ is also indicated to occur in the kidney^12, 80^ and has more recently been suggested to be involved in the metabolic events observed in diabetic kidney disease and ageing^81, 82^. Although the biochemistry of the Warburg effect is not fully understood, this phenomenon is consistent with our current observations.

NADH produced by the activation of glycolysis is not only used for lactate generation, but is also oxidized by the NADH shuttle (e.g., the malate-aspartate shuttle). The increased release of pyruvate into the renal vein after HS suggests that not all of the excess NADH in the cytoplasm produced by activation of the glycolytic system is used for lactate production. Enzymes of malate-aspartate shuttle have been identified in the kidney^83^ and gene expression of key enzymes in this pathway was observed in our study. The malate-aspartate shuttle has largely studied in cancer cells, and some suggest that the glucose fermentation (i.e. Warburg effect) is a secondary consequence of saturation of the shuttle^84^. The malate-aspartate shuttle can be stimulated by an increase in glutamine uptake ^85, 86^ which we observed (**Figure 7**). The key enzymes, oxaloacetate transaminase (GOT) and malate dehydrogenase (MDH) activate the shuttle by forming a complex with acetylation^87^ and we observed an increase in GOT mRNA expression. Given the recognized limitations of predicting the activity of the shuttle from gene expression levels, the data are consistent with the idea that the malate-aspartate shuttle is regenerating NADH inside of the mitochondrial matrix and sustaining oxidative phosphorylation.

### The high salt diet alters amino acid metabolism

The kidneys play a major role in the homeostasis of the body amino acid pools through the synthesis, degradation, filtration, reabsorption and urinary excretion of these compounds. Studies carried out in fasted swine found glutamine and proline from the arterial blood are largely disposed of by the kidneys and other amino acids such as serine, tyrosine and arginine generated and released from the kidneys for export to other tissues^17^. The current study carried out in unanesthetized non-fasted rats found glutamine was not taken up at LS state but a strong tendency for an increased uptake of glutamine (p=0.08) was observed during HS feeding. The kidneys also play an important role in protein metabolism which is filtered by the glomerulus^88^ is taken up into the lysosomes of the tubules and degraded to amino acids^89, 90^. Even the OM of the kidney appears to participate in amino acid metabolic function in the SD rat. Specifically, a clear reduction was observed in the metabolites related to the degradation of lysine in the OM at HS14 (**Figure S8B**) which then tended to return toward LS levels at HS21. This included reductions of α-ketoglutarate, glutamate, and allysine in the Lysine degradation pathway. Lysine has recently attracted attention for its ability to suppress salt-sensitive hypertension^91^. Although lysine is one of the essential amino acids, we found it was released from the SD rat kidney after HS (**Figure 7**), consistent with observations by Jang et al.^17^ in pig study. This is likely explained by the degradation of protein either by glomerular epithelial or tubular lysosomes during renal passage^92-94^. Moreover, we observed almost all the amino acids were released from kidney into renal vein in SD rats fed LS and further release was observed in most of detected amino acids when fed HS. In general, the data indicate that the kidney produces amino acids from protein degradation faster than their utilization by the kidney and that the HS diet enhanced proteolysis. There were, however, several notable exceptions to this such as glutamine which was released from the kidney at LS and tended to be taken up after HS. This uptake of glutamine may be involved in the activation of the malate-aspartate shuttle pathway as discussed earlier.

It was also found that the megalin (*Lrp2*) and clathrin (*Cltc*) mRNA expression levels were reduced with the HS diet (**Figure S6**) whereas several proteases and plasmid partitioning (PAR genes) were upregulated in Cx. Megalin and clathrin are key players in apical endocytosis in PT and reduction of these proteins is related to a reduction in albumin endocytosis^94, 95^. Megalin is downregulated with HS diets even in salt insensitive Wistar or SD rats and in the absence of increased urinary albumin excretion^96, 97^. However, as discussed earlier, the fact that the expression of transporter genes in the renal cortical proximal tubules appear to be decreased while the expression of genes distal to the TAL is increased suggests that proteolysis may be shifted to the distal tubules. For example, *Ctsa* and *Ctsb*, which are predominantly expressed in the proximal tubules, are downregulated, while *Ctsc* and *Ctsd*, which are highly expressed in the DCT and other areas^54^, are upregulated after salt loading. *Lrp2* and *Cltc* expressed in PT but not in DCT. It is known that DCT performs endocytosis of proteins, but it is not known which proteins play a key role, and further segment-specific studies are needed.

### Limitation

Several limitations must be recognized regarding the present study. First, rats were not fasted overnight in this study to avoid the known effects of fasting particularly on glucose/gluconeogenic metabolism. Although all samples were collected at the same time of day to avoid diurnal effects, the ad lib eating/drinking undoubtedly increase the variance of our results. Second, although RBF is conventionally reported in terms of volume flow per kidney weight, when RBF is continuously measured this normalization could not be applied since kidneys were weighted only at the end of the 21-day period of the HS diet. Based on other groups of SD rats that were studied, we observed no difference in kidney size between LS and rats fed HS for 21 days. RBF was therefore normalized by body weight recognizing that additional controls would need to be considered for studies in which the intervention produced significant changes of kidney weight. Third, despite the great progress that has been made in metabolite identification and the sensitivity of the mass spectrometry techniques, continuing efforts must be made in the development of the databases and informatics to link substrates, enzymes and metabolites to biochemical and physiological pathways. At the present time, much of this integration must be carried out manually. It is therefore important to carry out parallel transcriptomic or proteomic analysis since together these data can enhance the identification and validation of functional pathways. Fourth, the present study provides only directional changes in metabolite concentrations and it is evident that greater numbers of pure standards will be needed to enable large scale quantitative analysis.

### Conclusion

The kidneys of even normal SD rats with low blood pressure salt sensitivity exhibited significant changes in the metabolomic profiles in order to sustain the increased transport workloads and energy needs of the kidneys. The temporal patterns identified unique metabolic changes in the first 14 days of HS followed by what appear to compensatory response required to sustain the energy requirements of the kidney. It appears that the glycolysis was enhanced, and the production of pyruvate and lactate increased despite the increased oxygen consumption, while the TCA cycle was down regulated by the HS diet in SD rat’s kidney cortex. NADH produced during the process of increased glycolysis is used for lactate production in the cytoplasm and is involved in oxidative phosphorylation in mitochondria via malate-aspartate shuttle, which may contribute to oxygen consumption. Besides the activation of glycolysis, the oxidative pentose phosphate pathway, uncoupling protein 2 and nitric oxide synthetase 3 were upregulated each of which scavenge ROS and protect against kidney damage in SD rat’s kidney. The progressive increase of energy required for the high salt diet in face of a reduction in TCA cycle activity may be sustained by a “Warburg-like” effect whereby glucose and other six-carbon sugars are converted by glycolysis into cellular energy and the metabolite lactate which we found elevated in the renal venous blood (e.g., lactic acid fermentation). Although not previously identified in kidney cells, it is recognized that lactic acid fermentation can occur in muscle cells undergoing intense activity enabling ATP and NAD+ production to continue glycolysis ^98^. Finally, although kidney proteolysis appears to be enhanced by the HS diet, the metabolic consequences of this is unclear.

## Supporting information

mRNAseq method

## Acknowledgements

All metabolomics protocols were developed in and the mass spectrometry data was collected in the Mass Spectrometry and Protein Chemistry Lab within Protein Sciences at The Jackson Laboratory.

## References

1. Opie LH. Acute metabolic response in myocardial infarction. Heart. 1971;33:129–137.

2. Dulloo AG and Jacquet J. Adaptive reduction in basal metabolic rate in response to food deprivation in humans: a role for feedback signals from fat stores. The American journal of clinical nutrition. 1998;68:599–606.

3. Marosi K, Moehl K, Navas-Enamorado I, Mitchell SJ, Zhang Y, Lehrmann E, Aon MA, Cortassa S, Becker KG and Mattson MP. Metabolic and molecular framework for the enhancement of endurance by intermittent food deprivation. The FASEB Journal. 2018;32:3844–3858.

4. Weinberger MH. Salt Sensitivity of Blood Pressure in Humans. Hypertension. 1996;27:481–490.

5. Sanada H, Jones JE and Jose PA. Genetics of Salt-Sensitive Hypertension. Current Hypertension Reports. 2011;13:55–66.

6. Sullivan JM, Prewitt RL and Ratts TE. Sodium sensitivity in normotensive and borderline hypertensive humans. The American journal of the medical sciences. 1988;295:370–377.

7. Whittle JC. Does Racial Variation in Risk Factors Explain Black-White Differences in the Incidence of Hypertensive End-Stage Renal Disease? Archives of Internal Medicine. 1991;151:1359.

8. Klag MJ. End-stage Renal Disease in African-American and White Men. JAMA. 1997;277:1293.

9. Lipworth L, Mumma MT, Cavanaugh KL, Edwards TL, Ikizler TA, E. Tarone R, McLaughlin JK and Blot WJ. Incidence and Predictors of End Stage Renal Disease among Low-Income Blacks and Whites. PLoS ONE. 2012;7:e48407.

10. Vallis M. Sustained behaviour change in healthy eating to improve obesity outcomes: It is time to abandon willpower to appreciate wanting. Clinical Obesity. 2019;9:e12299.

11. Schultes B, Ernst B, Wilms B, Thurnheer M and Hallschmid M. Hedonic hunger is increased in severely obese patients and is reduced after gastric bypass surgery. The American journal of clinical nutrition. 2010;92:277–283.

12. Klahr S, Hamm LL, Hammerman MR and Mandel LJ. Renal Metabolism: Integrated Responses. Comprehensive Physiology. 2011.

13. Singh P, McDonough A, Thomson S, Skorecki K, Chertow G and Marsden P. Metabolic basis of solute transport. Brenner and Rector’s the kidney. 1. 2016.

14. Tian Z and Liang M. Renal metabolism and hypertension. Nature Communications. 2021;12.

15. Kim YJ and Felig P. Maternal and amniotic fluid substrate levels during caloric deprivation in human pregnancy. Metabolism. 1972;21:507–512.

16. Bing RJ. Cardiac metabolism. Physiological Reviews. 1965;45:171–213.

17. Jang C, Hui S, Zeng X, Cowan AJ, Wang L, Chen L, Morscher RJ, Reyes J, Frezza C, Hwang HY, Imai A, Saito Y, Okamoto K, Vaspoli C, Kasprenski L, Zsido GA, Gorman JH, Gorman RC and Rabinowitz JD. Metabolite Exchange between Mammalian Organs Quantified in Pigs. Cell Metabolism. 2019;30:594–606.e3.

18. Rinschen MM, Palygin O, Guijas C, Palermo A, Palacio-Escat N, Domingo-Almenara X, Montenegro-Burke R, Saez-Rodriguez J, Staruschenko A and Siuzdak G. Metabolic rewiring of the hypertensive kidney. Science signaling. 2019;12:eaax9760.

19. Kuehnbaum NL and Britz-Mckibbin P. New Advances in Separation Science for Metabolomics: Resolving Chemical Diversity in a Post-Genomic Era. Chemical Reviews. 2013;113:2437–2468.

20. Clish CB. Metabolomics: an emerging but powerful tool for precision medicine. Molecular Case Studies. 2015;1:a000588.

21. Murray GC, Bais P, Hatton CL, Tadenev ALD, Hoffmann BR, Stodola TJ, Morelli KH, Pratt SL, Schroeder D, Doty R, Fiehn O, John SWM, Bult CJ, Cox GA and Burgess RW. Mouse models of NADK2 deficiency analyzed for metabolic and gene expression changes to elucidate pathophysiology. Human Molecular Genetics. 2022.

22. Shimada S, Yang C, Kurth T and Cowley Jr AW. Divergent roles of angiotensin II upon the immediate and sustained increases of renal blood flow following unilateral nephrectomy. American Journal of Physiology-Renal Physiology. 2022;322:F473–F485.

23. Shimada S and Cowley Jr AW. Long-Term Continuous Measurement of Renal Blood Flow in Conscious Rats. Journal of Visualized Experiments: Jove. 2022.

24. Mori T and Cowley Jr AW. Role of pressure in angiotensin II-induced renal injury: chronic servo-control of renal perfusion pressure in rats. Hypertension. 2004;43:752–759.

25. Evans LC, Petrova G, Kurth T, Yang C, Bukowy JD, Mattson DL and Cowley Jr AW. Increased perfusion pressure drives renal T-cell infiltration in the Dahl salt-sensitive rat. Hypertension. 2017;70:543–551.

26. Shimada S, Abais-Battad JM, Alsheikh AJ, Yang C, Stumpf M, Kurth T, Mattson DL and Cowley Jr AW. Renal Perfusion Pressure Determines Infiltration of Leukocytes in the Kidney of Rats With Angiotensin II–Induced Hypertension. Hypertension. 2020;76:849–858.

27. Evans LC, Ryan RP, Broadway E, Skelton MM, Kurth T and Cowley AW. Null Mutation of the Nicotinamide Adenine Dinucleotide Phosphate–Oxidase Subunit p67phox Protects the Dahl-S Rat From Salt-Induced Reductions in Medullary Blood Flow. Hypertension. 2015;65:561–568.

28. Cowley AW, Ryan RP, Kurth T, Skelton MM, Schock-Kusch D and Gretz N. Progression of Glomerular Filtration Rate Reduction Determined in Conscious Dahl Salt-Sensitive Hypertensive Rats. Hypertension. 2013;62:85–90.

29. Friedemann J, Heinrich R, Shulhevich Y, Raedle M, William-Olsson L, Pill J and Schock-Kusch D. Improved kinetic model for the transcutaneous measurement of glomerular filtration rate in experimental animals. Kidney International. 2016;90:1377–1385.

30. Welch WJ, Baumgärtl H, Lübbers D and Wilcox CS. Nephron pO2 and renal oxygen usage in the hypertensive rat kidney. Kidney International. 2001;59:230–237.

31. Mik EG, Johannes T and Ince C. Monitoring of renal venous PO2 and kidney oxygen consumption in rats by a near-infrared phosphorescence lifetime technique. American Journal of Physiology-Renal Physiology. 2008;294:F676–F681.

32. Xia J, Psychogios N, Young N and Wishart DS. MetaboAnalyst: a web server for metabolomic data analysis and interpretation. Nucleic acids research. 2009;37:W652–W660.

33. Cantrell CD. Modern mathematical methods for physicists and engineers: Cambridge University Press; 2000.

34. Wu Y-L, Hu C-Y, Lu S-S, Gong F-F, Feng F, Qian Z-Z, Ding X-X, Yang H-Y and Sun Y-H. Association between methylenetetrahydrofolate reductase (MTHFR) C677T/A1298C polymorphisms and essential hypertension: a systematic review and meta-analysis. Metabolism. 2014;63:1503–1511.

35. Seufert L, Benzing T, Ignarski M and Müller R-U. RNA-binding proteins and their role in kidney disease. Nature Reviews Nephrology. 2022;18:153–170.

36. Brooks GA. Cell–cell and intracellular lactate shuttles. The Journal of physiology. 2009;587:5591–5600.

37. Bergmann M. A classification of proteolytic enzymes. Advances in Enzymology (eds FF Nord, CH Werkman). 2006;2:49–67.

38. Nielsen R, Mollet G, Esquivel EL, Weyer K, Nielsen PK, Antignac C and Christensen EI. Increased lysosomal proteolysis counteracts protein accumulation in the proximal tubule during focal segmental glomerulosclerosis. Kidney International. 2013;84:902–910.

39. Palygin O, Ilatovskaya DV and Staruschenko A. Protease-activated receptors in kidney disease progression. American Journal of Physiology-Renal Physiology. 2016;311:F1140–F1144.

40. Roman RJ and Fan F. 20-HETE. Hypertension. 2018;72:12–18.

41. Allison PD. Fixed effects regression models: SAGE publications; 2009.

42. Morimoto A, Uzu T, Fujii T, Nishimura M, Kuroda S, Nakamura S, Inenaga T and Kimura G. Sodium sensitivity and cardiovascular events in patients with essential hypertension. The Lancet. 1997;350:1734–1737.

43. Elijovich F, Weinberger MH, Anderson CAM, Appel LJ, Bursztyn M, Cook NR, Dart RA, Newton-Cheh CH, Sacks FM and Laffer CL. Salt Sensitivity of Blood Pressure. Hypertension. 2016;68:e7–e46.

44. Weinberger MH, Fineberg NS, Fineberg SE and Weinberger M. Salt Sensitivity, Pulse Pressure, and Death in Normal and Hypertensive Humans. Hypertension. 2001;37:429–432.

45. Isaksson GL, Stubbe J, Lyngs Hansen P, Jensen BL and Bie P. Salt sensitivity of renin secretion, glomerular filtration rate and blood pressure in conscious Sprague-Dawley rats. Acta Physiologica. 2014;210:446–454.

46. Mose FH, Jörgensen AN, Vrist MH, Ekelöf NP, Pedersen EB and Bech JN. Effect of 3% saline and furosemide on biomarkers of kidney injury and renal tubular function and GFR in healthy subjects – a randomized controlled trial. BMC Nephrology. 2019;20.

47. Parmer RJ, Stone RA and Cervenka JH. Renal hemodynamics in essential hypertension. Racial differences in response to changes in dietary sodium. Hypertension. 1994;24:752–757.

48. Koomans HA, Roos JC, Dorhout Mees EJ and Delawi IM. Sodium balance in renal failure. A comparison of patients with normal subjects under extremes of sodium intake. Hypertension. 1985;7:714–721.

49. Pechère-Bertschi A, Maillard M, Stalder H, Bischof P, Fathi M, Brunner HR and Burnier M. Renal hemodynamic and tubular responses to salt in women using oral contraceptives. Kidney International. 2003;64:1374–1380.

50. Weir MR, Dengel DR, Behrens MT and Goldberg AP. Salt-induced increases in systolic blood pressure affect renal hemodynamics and proteinuria. Hypertension. 1995;25:1339–1344.

51. Udwan K, Abed A, Roth I, Dizin E, Maillard M, Bettoni C, Loffing J, Wagner CA, Edwards A and Feraille E. Dietary sodium induces a redistribution of the tubular metabolic workload. The Journal of Physiology. 2017;595:6905–6922.

52. Hansell P, Welch WJ, Blantz RC and Palm F. Determinants of kidney oxygen consumption and their relationship to tissue oxygen tension in diabetes and hypertension. Clinical and Experimental Pharmacology and Physiology. 2013;40:123–137.

53. Kiil F, Aukland K and Refsum HE. Renal sodium transport and oxygen consumption. American Journal of Physiology-Legacy Content. 1961;201:511–516.

54. Lee JW, Chou C-L and Knepper MA. Deep Sequencing in Microdissected Renal Tubules Identifies Nephron Segment–Specific Transcriptomes. Journal of the American Society of Nephrology. 2015;26:2669–2677.

55. Pavlov TS, Levchenko V, O’Connor PM, Ilatovskaya DV, Palygin O, Mori T, Mattson DL, Sorokin A, Lombard JH and Cowley AW. Deficiency of renal cortical EGF increases ENaC activity and contributes to salt-sensitive hypertension. Journal of the American Society of Nephrology. 2013;24:1053–1062.

56. Blass G, Klemens CA, Brands MW, Palygin O and Staruschenko A. Postprandial effects on ENaC-mediated sodium absorption. Scientific reports. 2019;9:1–11.

57. Spencer NY and Stanton RC. Glucose 6-phosphate dehydrogenase and the kidney. Current opinion in nephrology and hypertension. 2017;26:43–49.

58. Aguilar E, Esteves P, Sancerni T, Lenoir V, Aparicio T, Bouillaud F, Dentin R, Prip-Buus C, Ricquier D, Pecqueur C, Guilmeau S and Alves-Guerra M-C. UCP2 Deficiency Increases Colon Tumorigenesis by Promoting Lipid Synthesis and Depleting NADPH for Antioxidant Defenses. Cell Reports. 2019;28:2306–2316.e5.

59. Cowley Jr AW, Mori T, Mattson D and Zou A-P. Role of renal NO production in the regulation of medullary blood flow. American Journal of Physiology-Regulatory, Integrative and Comparative Physiology. 2003;284:R1355–R1369.

60. Abais-Battad JM, Saravia FL, Lund H, Dasinger JH, Fehrenbach DJ, Alsheikh AJ, Zemaj J, Kirby JR and Mattson DL. Dietary influences on the Dahl SS rat gut microbiota and its effects on salt-sensitive hypertension and renal damage. Acta Physiologica. 2021;232:e13662.

61. Abais-Battad JM and Mattson DL. Influence of dietary protein on Dahl salt-sensitive hypertension: a potential role for gut microbiota. American Journal of Physiology-Regulatory, Integrative and Comparative Physiology. 2018;315:R907–R914.

62. Kato T, Niizuma S, Inuzuka Y, Kawashima T, Okuda J, Kawamoto A, Tamaki Y, Iwanaga Y, Soga T and Kita T. Analysis of liver metabolism in a rat model of heart failure. International journal of cardiology. 2012;161:130–136.

63. Welch WJ and Wilcox CS. Modulating role for thromboxane in the tubuloglomerular feedback response in the rat. Journal of Clinical Investigation. 1988;81:1843–1849.

64. Roman RJ. P-450 metabolites of arachidonic acid in the control of cardiovascular function. Physiological reviews. 2002;82:131–185.

65. Yamashita W, Ito Y, Weiss MA, Ooi BS and Pollak VE. A thromboxane synthetase antagonist ameliorates progressive renal disease of Dahl-S rats. Kidney international. 1988;33:77–83.

66. Fujii K, Onaka U, Ohya Y, Ohmori S, Tominaga M, Abe I, Takata Y and Fujishima M. Role of eicosanoids in alteration of membrane electrical properties in isolated mesenteric arteries of salt-loaded, Dahl salt-sensitive rats. British Journal of Pharmacology. 1997;120:1207–1214.

67. Ren Y, D’Ambrosio MA, Garvin JL, Peterson EL and Carretero OA. Mechanism of impaired afferent arteriole myogenic response in Dahl salt-sensitive rats: role of 20-HETE. American Journal of Physiology-Renal Physiology. 2014;307:F533–F538.

68. Uehara Y, Tobian L and Iwai J. Platelet thromboxane inhibition by plasma polypeptides in prehypertensive Dahl rats. Hypertension. 1986;8:II180.

69. Salvati P, Ferti C, Ferrario RG, Lamberti E, Duzzi L, Bianchi G, Remuzzi G, Perico N, Benigni A and Braidotti P. Role of enhanced glomerular synthesis of thromboxane A2 in progressive kidney disease. Kidney international. 1990;38:447–458.

70. Wang T, Fu X, Chen Q, Patra JK, Wang D, Wang Z and Gai Z. Arachidonic Acid Metabolism and Kidney Inflammation. International Journal of Molecular Sciences. 2019;20:3683.

71. Schaub JA, Venkatachalam MA and Weinberg JM. Proximal tubular oxidative metabolism in acute kidney injury and the transition to CKD. Kidney360. 2021;2:355.

72. Wang Y, Liu X, Zhang C and Wang Z. High salt diet induces metabolic alterations in multiple biological processes of Dahl salt-sensitive rats. The Journal of nutritional biochemistry. 2018;56:133–141.

73. Tanada Y, Okuda J, Kato T, Minamino-Muta E, Murata I, Soga T, Shioi T and Kimura T. The metabolic profile of a rat model of chronic kidney disease. PeerJ. 2017;5:e3352.

74. Zheleznova NN, Yang C, Ryan RP, Halligan BD, Liang M, Greene AS and Cowley Jr AW. Mitochondrial proteomic analysis reveals deficiencies in oxygen utilization in medullary thick ascending limb of Henle in the Dahl salt-sensitive rat. Physiological genomics. 2012.

75. Potter JC, Whiles SA, Miles CB, Whiles JB, Mitchell MA, Biederman BE, Dawoud FM, Breuel KF, Williamson GA, Picken MM and Polichnowski AJ. Salt-Sensitive Hypertension, Renal Injury, and Renal Vasodysfunction Associated With Dahl Salt-Sensitive Rats Are Abolished in Consomic SS.BN1 Rats. Journal of the American Heart Association. 2021;10.

76. Makino A, Skelton MM, Zou A-P and Cowley AW. Increased Renal Medullary H2O2 Leads to Hypertension. Hypertension. 2003;42:25–30.

77. Makino A, Skelton MM, Zou A-P, Roman RJ and Cowley AW. Increased Renal Medullary Oxidative Stress Produces Hypertension. Hypertension. 2002;39:667–672.

78. Miyata N and Cowley AW. Renal Intramedullary Infusion of l-Arginine Prevents Reduction of Medullary Blood Flow and Hypertension in Dahl Salt-Sensitive Rats. Hypertension. 1999;33:446–450.

79. Warburg O. The metabolism of carcinoma cells. The Journal of Cancer Research. 1925;9:148–163.

80. Lee JB and Peterhm. Effect of oxygen tension on glucose metabolism in rabbit kidney cortex and medulla. American Journal of Physiology-Legacy Content. 1969;217:1464–1471.

81. Zhang G, Darshi M and Sharma K. The Warburg effect in diabetic kidney disease. Seminars in nephrology. 2018;38:111–120.

82. Burns JS and Manda G. Metabolic pathways of the Warburg effect in health and disease: perspectives of choice, chain or chance. International journal of molecular sciences. 2017;18:2755.

83. Guder WG and Ross BD. Enzyme distribution along the nephron. Kidney international. 1984;26:101–111.

84. Wang Y, Stancliffe E, Fowle-Grider R, Wang R, Wang C, Schwaiger-Haber M, Shriver LP and Patti GJ. Saturation of the mitochondrial NADH shuttles drives aerobic glycolysis in proliferating cells. Molecular cell. 2022;82:3270–3283. e9.

85. Friedrichs D. On the stimulation of gluconeogenesis by L-lysine in isolated rat kidney cortex tubules. Biochimica et Biophysica Acta (BBA)-General Subjects. 1975;392:255–270.

86. Stumvoll M, Perriello G, Meyer C and Gerich J. Role of glutamine in human carbohydrate metabolism in kidney and other tissues. Kidney International. 1999;55:778–792.

87. Yang H, Zhou L, Shi Q, Zhao Y, Lin H, Zhang M, Zhao S, Yang Y, Ling ZQ, Guan KL, Xiong Y and Ye D. SIRT 3-dependent GOT2 acetylation status affects the malate–aspartate NADH shuttle activity and pancreatic tumor growth. The EMBO Journal. 2015;34:1110–1125.

88. Dickson LE, Wagner MC, Sandoval RM and Molitoris BA. The proximal tubule and albuminuria: really! Journal of the American Society of Nephrology. 2014;25:443–453.

89. Tojo A and Kinugasa S. Mechanisms of Glomerular Albumin Filtration and Tubular Reabsorption. International Journal of Nephrology. 2012;2012:1–9.

90. Mori KP, Yokoi H, Kasahara M, Imamaki H, Ishii A, Kuwabara T, Koga K, Kato Y, Toda N, Ohno S, Kuwahara K, Endo T, Nakao K, Yanagita M, Mukoyama M and Mori K. Increase of Total Nephron Albumin Filtration and Reabsorption in Diabetic Nephropathy. Journal of the American Society of Nephrology. 2017;28:278–289.

91. Rinschen MM, Palygin O, El-Meanawy A, Domingo-Almenara X, Palermo A, Dissanayake LV, Golosova D, Schafroth MA, Guijas C, Demir F, Jaegers J, Gliozzi ML, Xue J, Hoehne M, Benzing T, Kok BP, Saez E, Bleich M, Himmerkus N, Weisz OA, Cravatt BF, Krüger M, Benton HP, Siuzdak G and Staruschenko A. Accelerated lysine metabolism conveys kidney protection in salt-sensitive hypertension. Nature Communications. 2022;13.

92. Russo LM, Bakris GL and Comper WD. Renal handling of albumin: a critical review of basic concepts and perspective. American journal of kidney diseases. 2002;39:899–919.

93. Osicka TM and Comper WD. Protein degradation during renal passage in normal kidneys is inhibited in experimental albuminuria. Clinical science (London, England: 1979). 1997;93:65–72.

94. Russo L, Sandoval R, McKee M, Osicka T, Collins A, Brown D, Molitoris B and Comper W. The normal kidney filters nephrotic levels of albumin retrieved by proximal tubule cells: retrieval is disrupted in nephrotic states. Kidney international. 2007;71:504–513.

95. Christensen EI and Birn H. Megalin and cubilin: multifunctional endocytic receptors. Nature Reviews Molecular Cell Biology. 2002;3:258–267.

96. Washino S, Hosohata K, Jin D, Takai S and Miyagawa T. Early urinary biomarkers of renal tubular damage by a high-salt intake independent of blood pressure in normotensive rats. Clinical and Experimental Pharmacology and Physiology. 2018;45:261–268.

97. Jo SM, Nam J, Park S-Y, Park G, Kim BG, Jeong G-H, Hurh BS and Kim JY. Effect of Mineral-Balanced Deep-Sea Water on Kidney Function and Renal Oxidative Stress Markers in Rats Fed a High-Salt Diet. International Journal of Molecular Sciences. 2021;22:13415.

98. Manoj KM, Nirusimhan V, Parashar A, Edward J and Gideon DA. Murburn precepts for lactic-acidosis, Cori cycle, and Warburg effect: Interactive dynamics of dehydrogenases, protons, and oxygen. Journal of Cellular Physiology. 2022;237:1902–1922.

99. Berg JM, Tymoczko JL and Stryer L. Biochemistry. 7th ed. ed. New York: W.H. Freeman; 2012.

