## Supplemental information for "Metabolic responses of normal rat kidneys to a high salt intake"

**Authors and affiliations:**

Satoshi Shimada[1], Brian R. Hoffmann[2], Chun Yang[1], Theresa Kurth[1], Andrew S. Greene[2], Mingyu Liang [1], Ranjan K. Dash[1,3], Allen W. Cowley Jr[1] *.

1. Department of Physiology, Medical College of Wisconsin, Milwaukee, Wisconsin, USA.
2. Mass Spectrometry and Protein Chemistry, Protein Sciences, The Jackson Laboratory, Bar Harbor, Maine, USA
3. Department of Biomedical Engineering, Medical College of Wisconsin and Marquette University, Milwaukee, Wisconsin, USA.

*Corresponding author

Allen W. Cowley, Jr., Ph.D.

Department of Physiology, Medical College of Wisconsin

8701 Watertown Plank Rd, Milwaukee, WI 53226, USA

Figure S1.

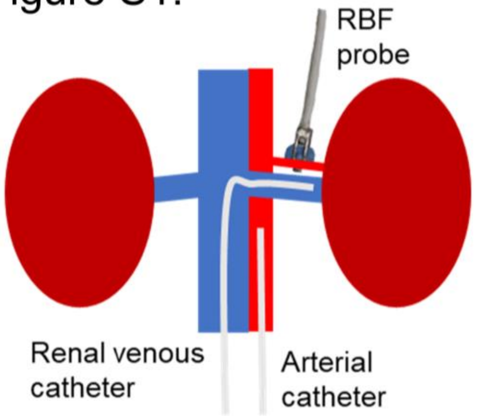

**Figure S1**. The schema of the chronically instrumented rat. Illustrated is the ultrasonic flow probe on the left renal artery to measure renal blood flow (RBF) and the chronically implanted aortic and renal venous catheters for intermittent sampling of blood.

Figure S2.

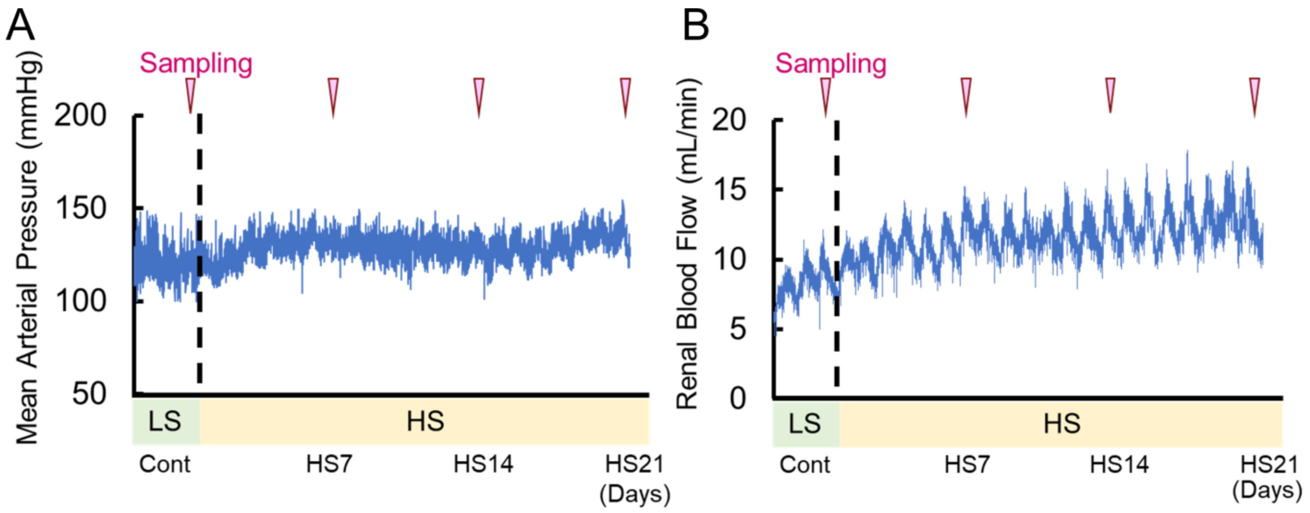

**Figure S2**. Representative (n=1) 1 min average of (A) mean arterial pressure and (B) renal blood flow, and periods of sampling for bloods and urine.

Figure S3.

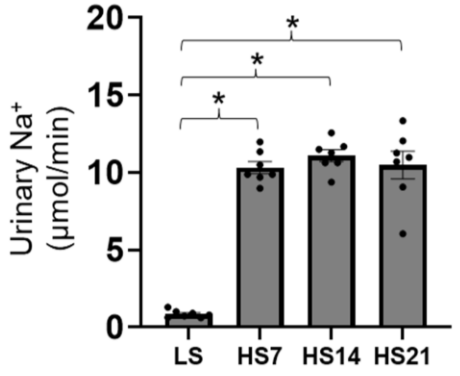

**Figure S3**. Urinary $Na^+$ excretion rates (n=7 rats) at each of the days of sampling (LS, HS7, HS14, and HS21. Mean ± SEM and individual data. *p<0.05 vs LS, One-way RM ANOVA, Holm-Sidak.

Figure S4.

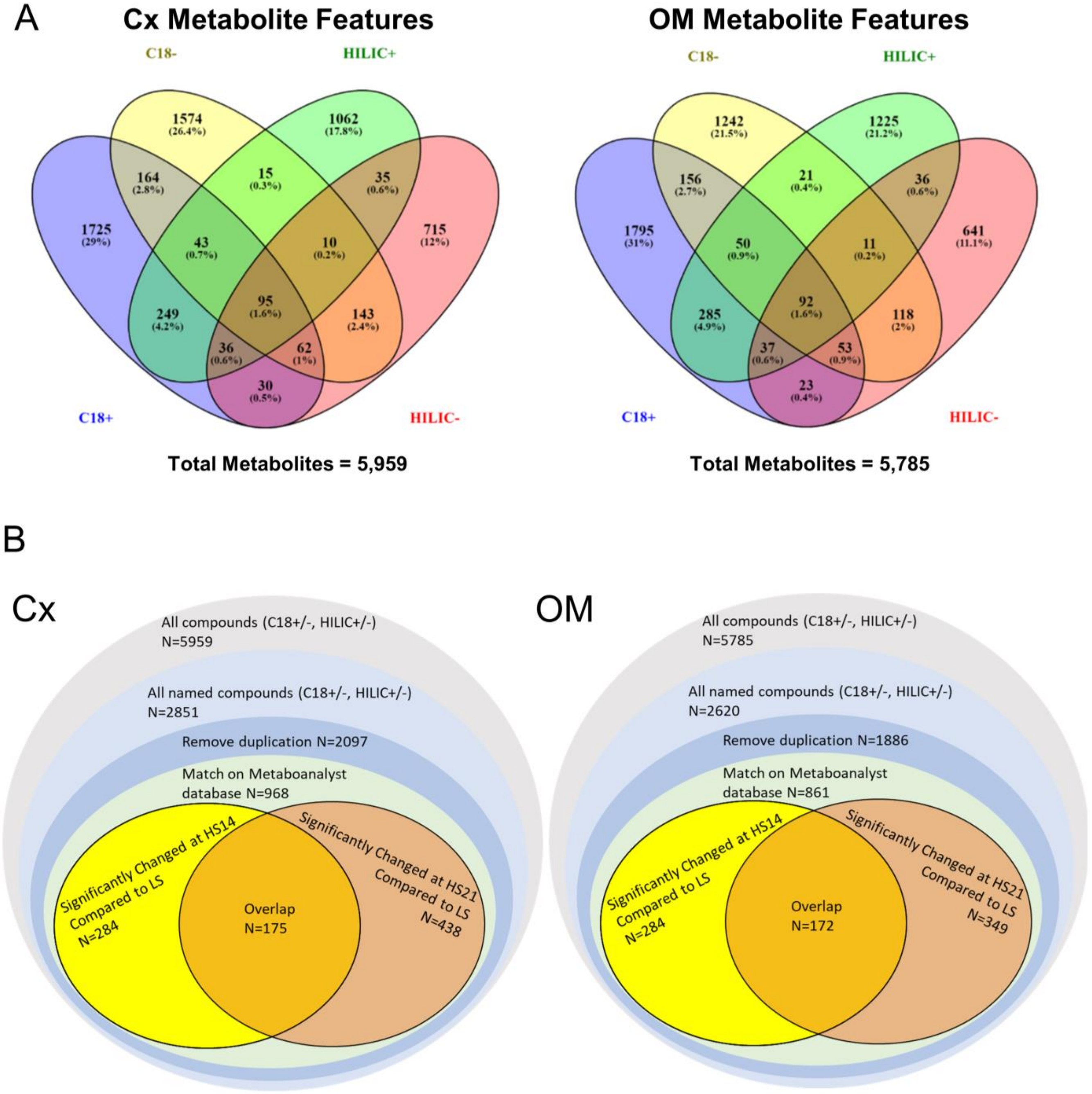

**Figure S4**. Venn diagrams of cortical (Cx) and outer medullary (OM) tissue metabolite features showing the number of metabolites detected in each of 4 modes (C18+/-, HILIC+/-). (A) Illustrates the number of metabolites detected, named and listed in the Metaboanalyst 5.0 database (October 2022). (B). Illustrates the number of those that significantly ($p<0.05$) differed from LS at HS by t-test.

### Figure S5.

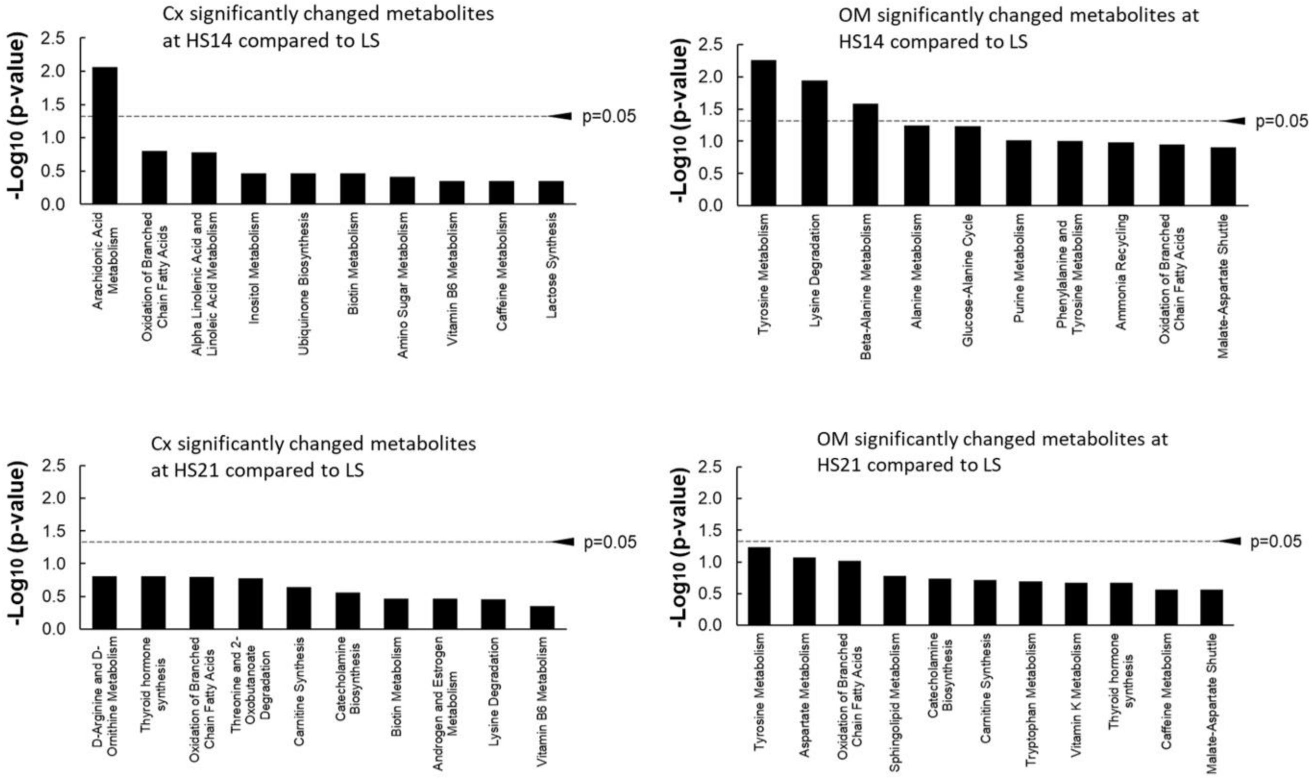

**Figure S5**. Results of the top 10 pathways identified in the cortex (Cx) and outer medulla (OM) from the metabolomic analysis determined by enrichment analysis on Metaboanalyst 5.0 (SMPDB, October 2022). Shown are pathways that were changed at HS days 14 and 21 compared to LS fed rats. The $-\log_{10}$ p-values are plotted with those above the dotted horizontal line representing $p<0.05$.

Figure S6.

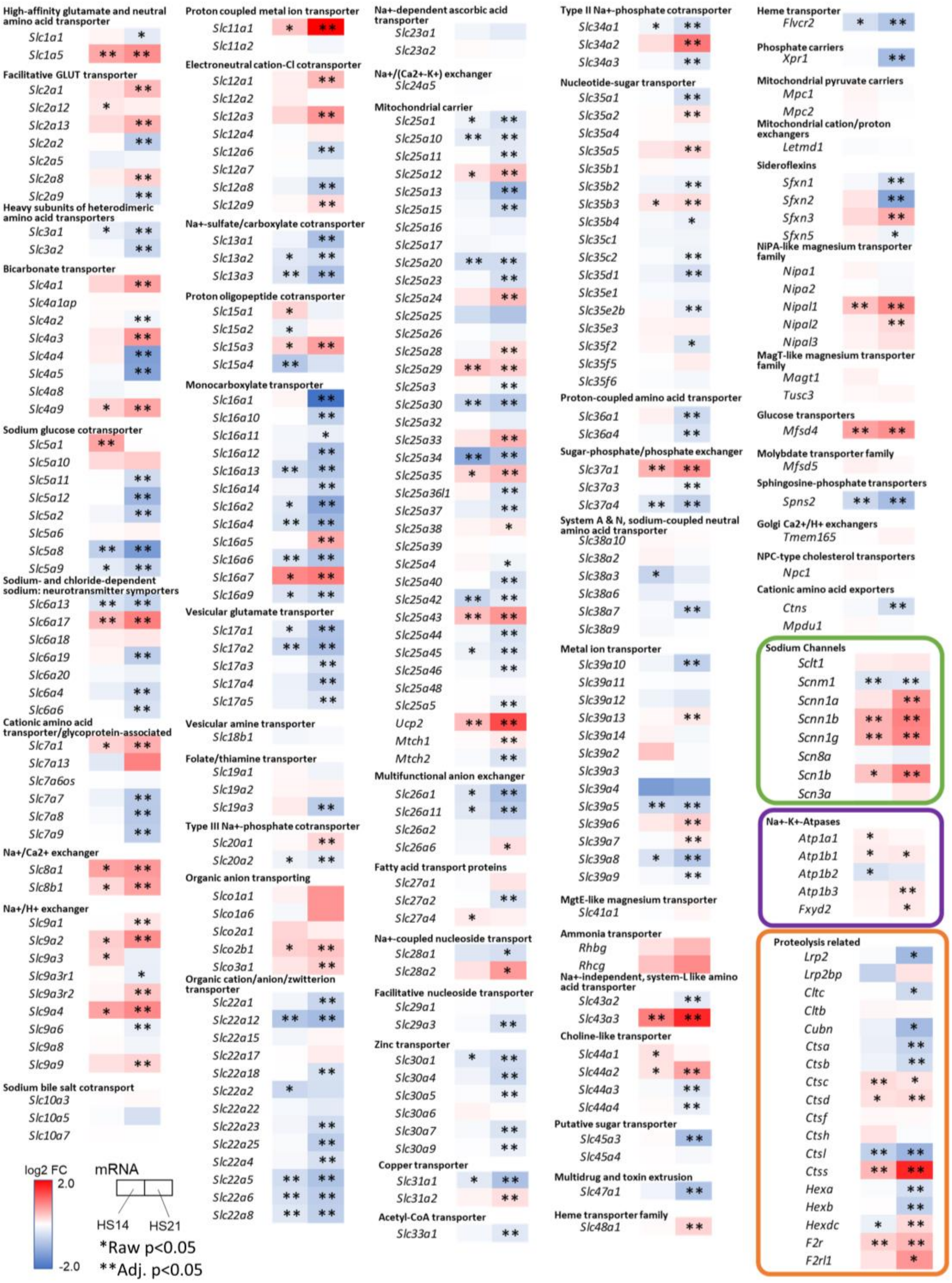

**Figure S6**. mRNA expression of transporters, channels, Na+-K+-ATP+ases and proteolysis of renal cortex (Cx) comparing $log_2$ fold change (FC) of high salt (HS) days 14 and 21 to low salt (LS). Red denotes increase in expression and blue denotes decrease in expression. Not framed are solute carrier family genes, framed in green are sodium channels, framed in purple are Na+-K+-ATP+ases and framed in orange are proteolysis related genes. *Raw p<0.05 in DESeq2, **Adj. p<0.05 in Benjamini and Hochberg.

Figure S7.

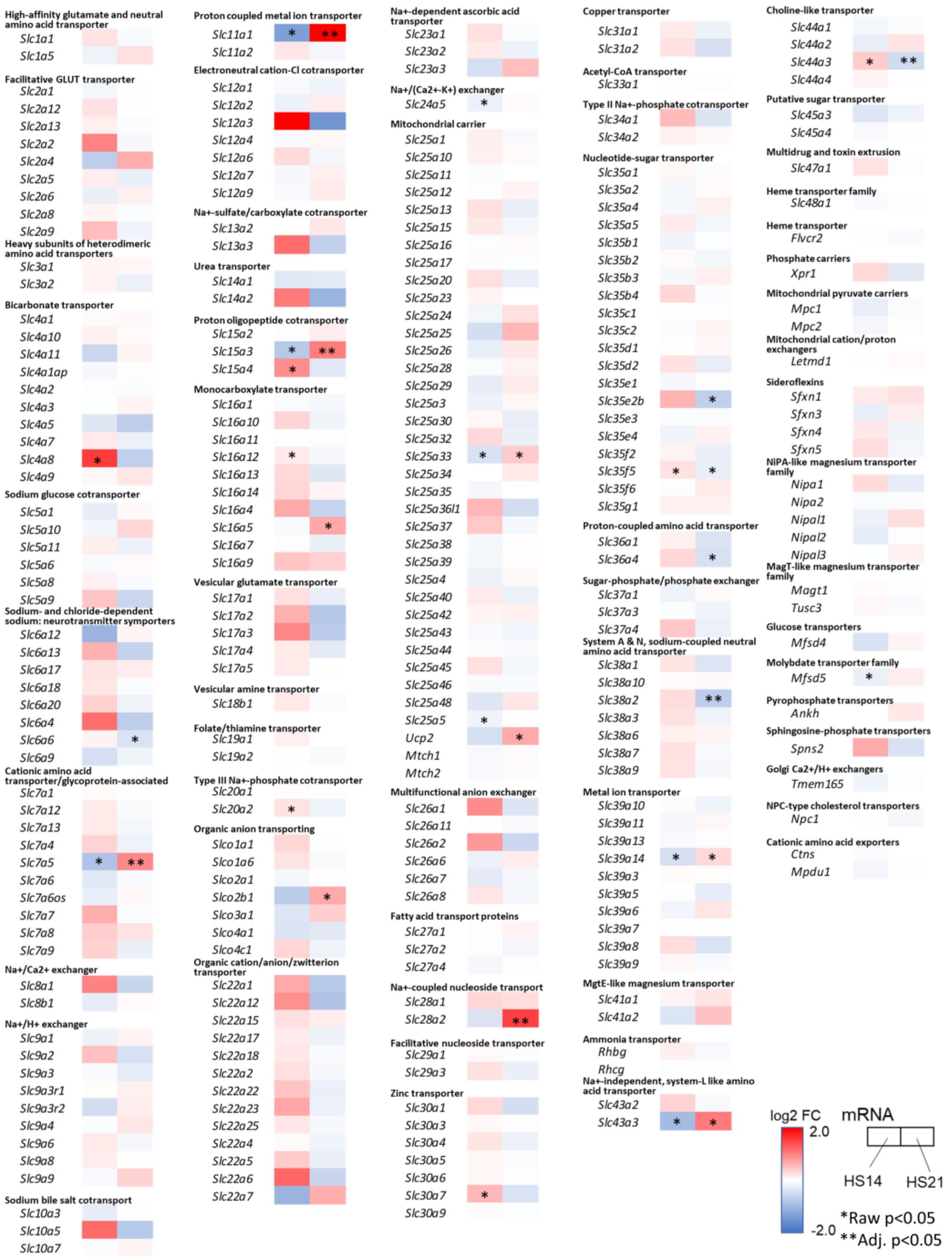

**Figure S7**. mRNA expression of transporter in outer medulla (OM).

Log$_2$ fold change (FC) of high salt (HS) to low salt (LS) are represented in color. The left boxes are FC of HS14 to LS and the right boxes are FC of HS21 to LS. Red denotes increase in expression and blue denotes decrease in expression *Raw p<0.05 in DESeq2, **Adj. p<0.05 in Benjamini and Hochberg.

Figure S8

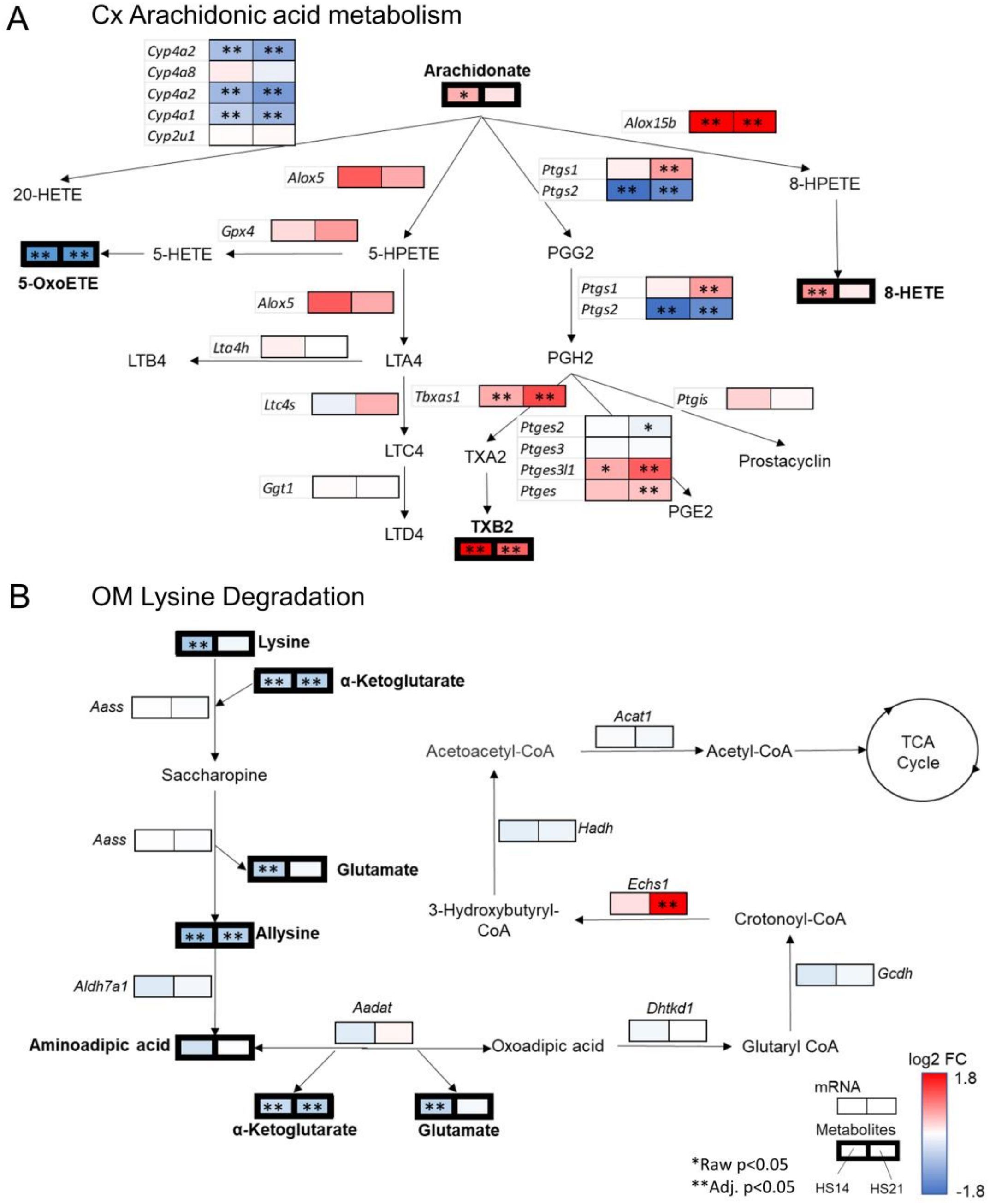

**Figure S8**. Integrated figure of arachidonic acid metabolism in cortex (Cx) (A) and lysine metabolism in outer medulla (OM) (B).

$Log_2$ fold change (FC) of high salt (HS) to low salt (LS) are represented in color. The left boxes are FC of HS14 to LS and the right boxes are FC of HS21 to LS. Thin boxes represent mRNA and thick boxes represent metabolites. Red denotes increase in expression and blue denotes decrease in expression. *Raw p<0.05 in t-test for metabolomics and in DESeq2 for mRNAseq, **Adj. p<0.05 in Benjamini and Hochberg. (KEGG map ID 00590 last update: September/28/2022, SMP0000037 last update: November/24/2022)

Figure S9.

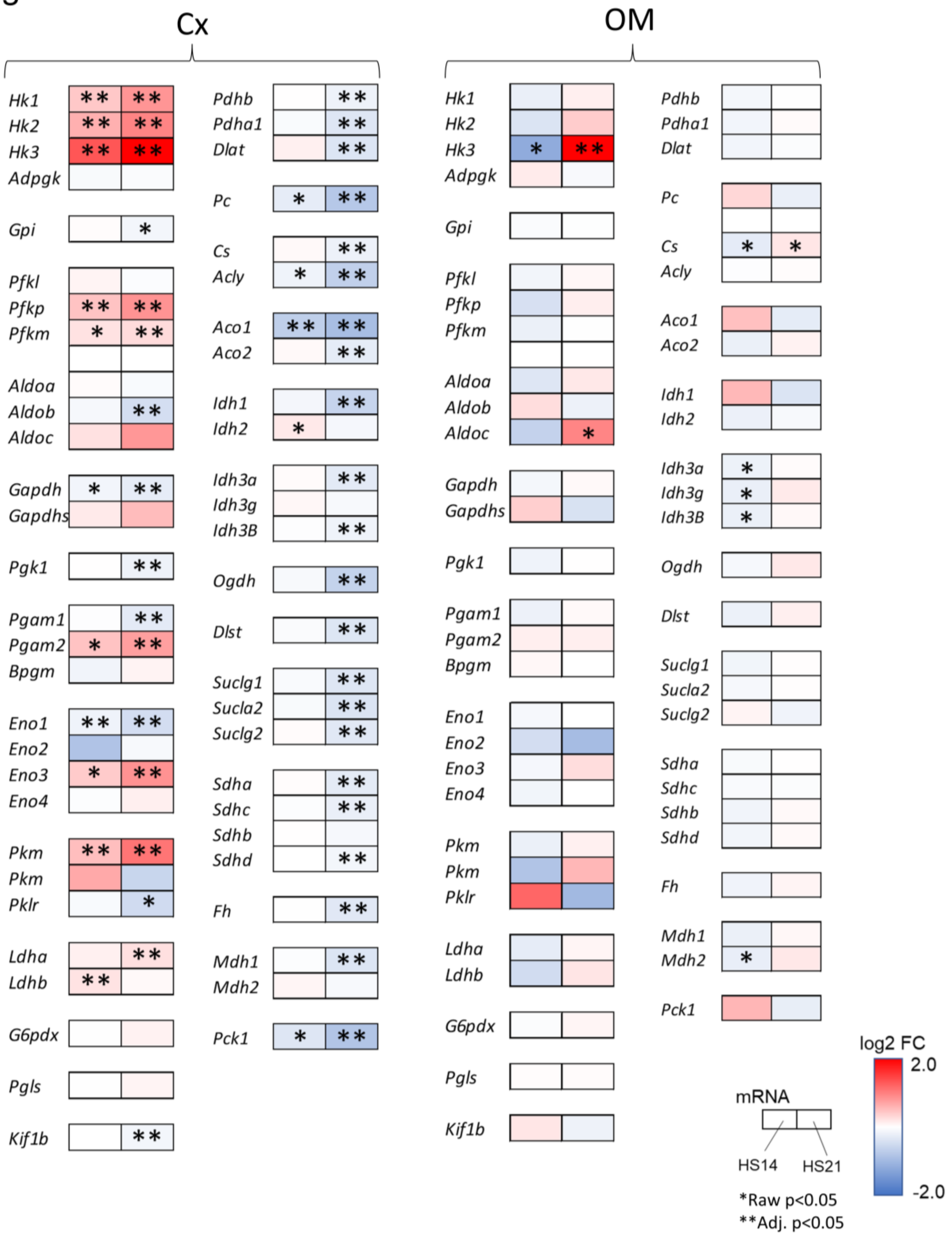

**Figure S9**. Gene expression of glycolysis and TCA cycle in cortex (Cx) and outer medulla (OM) $Log_2$ fold change (FC) of high salt (HS) to low salt (LS) are represented in color. The left boxes are FC of HS14 to LS and the right boxes are FC of HS21 to LS. Red denotes increase in expression and blue denotes decrease in expression *Raw p<0.05 in DESeq2, **Adj. p<0.05 in Benjamini and Hochberg.

Figure S10.

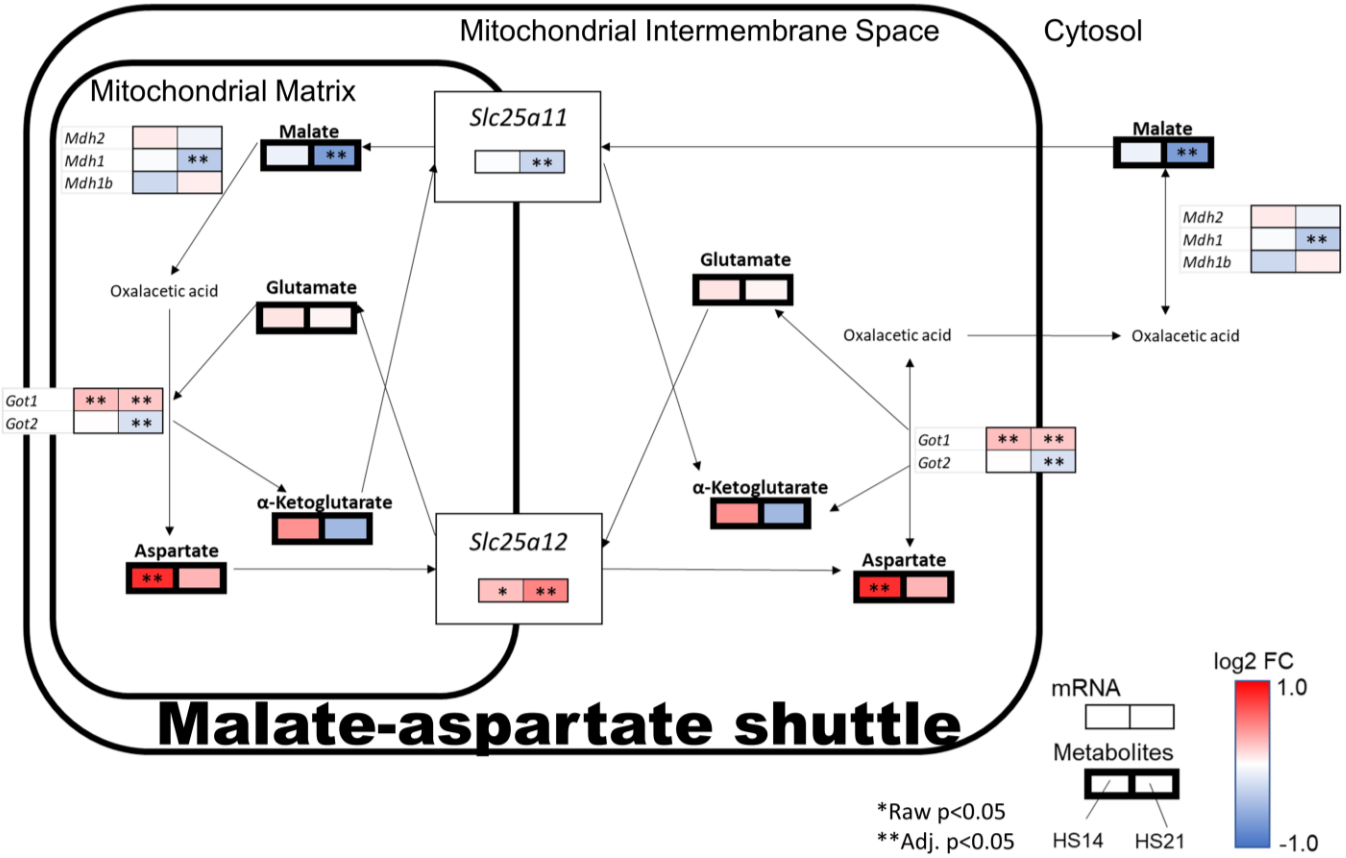

**Figure S10**. Integrated figure of malate-aspartate shuttle in cortex

$Log_2$ fold change (FC) of high salt (HS) to low salt (LS) are represented in color. The left boxes are FC of HS14 to LS and the right boxes are FC of HS21 to LS. Thin boxes represent mRNA and thick boxes represent metabolites. Red denotes increase in expression and blue denotes decrease in expression. *Raw p<0.05 in t-test for metabolomics and in DESeq2 for mRNAseq, **Adj. p<0.05 in Benjamini and Hochberg. (SMP0000129 last update: October/18/2022)

Figure S11.

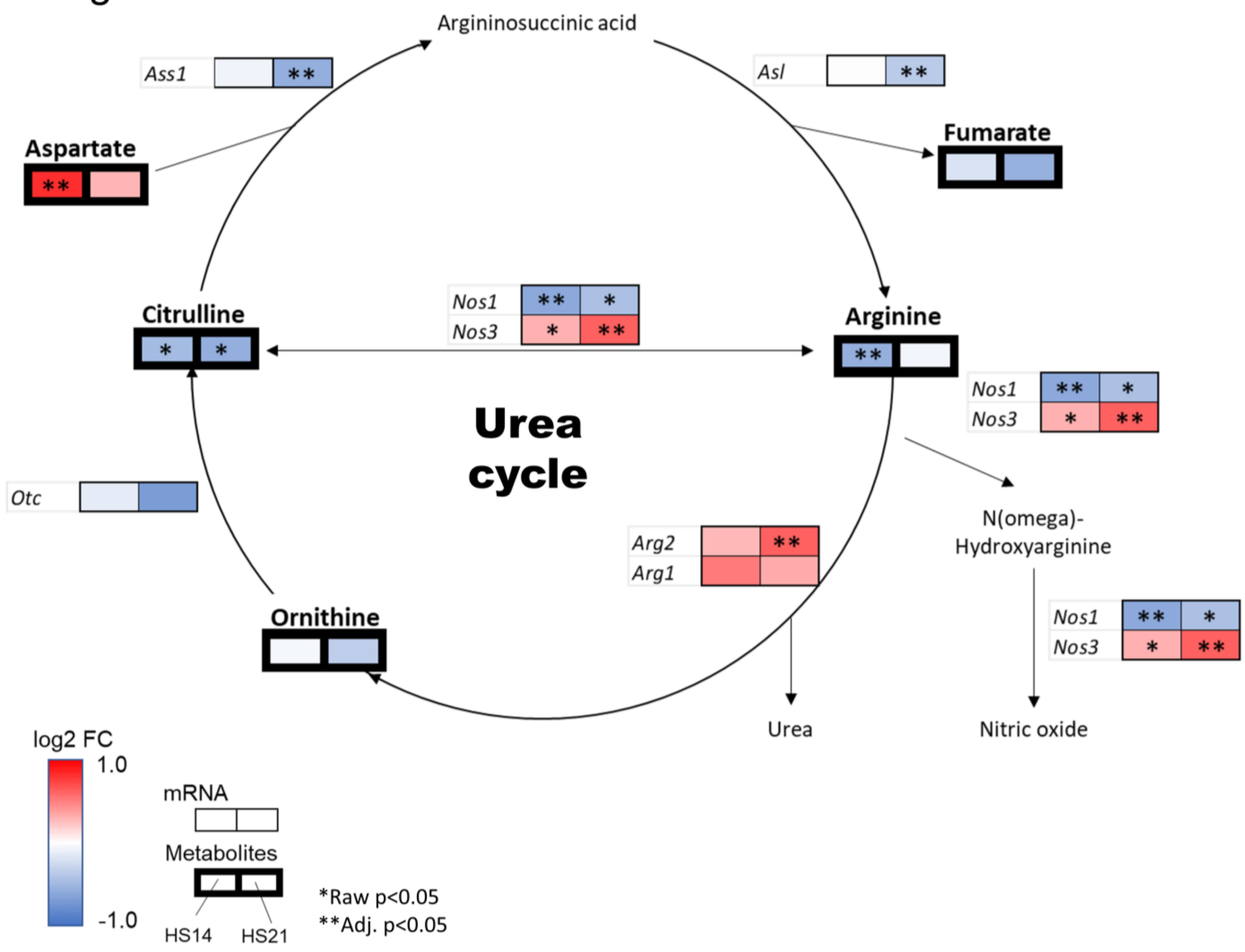

**Figure S11**. Integrated figure of urea cycle in cortex

$Log_2$ fold change (FC) of high salt (HS) to low salt (LS) are represented in color. The left boxes are FC of HS14 to LS and the right boxes are FC of HS21 to LS. Thin boxes represent mRNA and thick boxes represent metabolites. Red denotes increase in expression and blue denotes decrease in expression. *Raw p<0.05 in t-test for metabolomics and in DESeq2 for mRNAseq, **Adj. p<0.05 in Benjamini and Hochberg. (KEGG map ID 00220 last update: July/29/2022)

Figure S12. Metabolite features in plasma

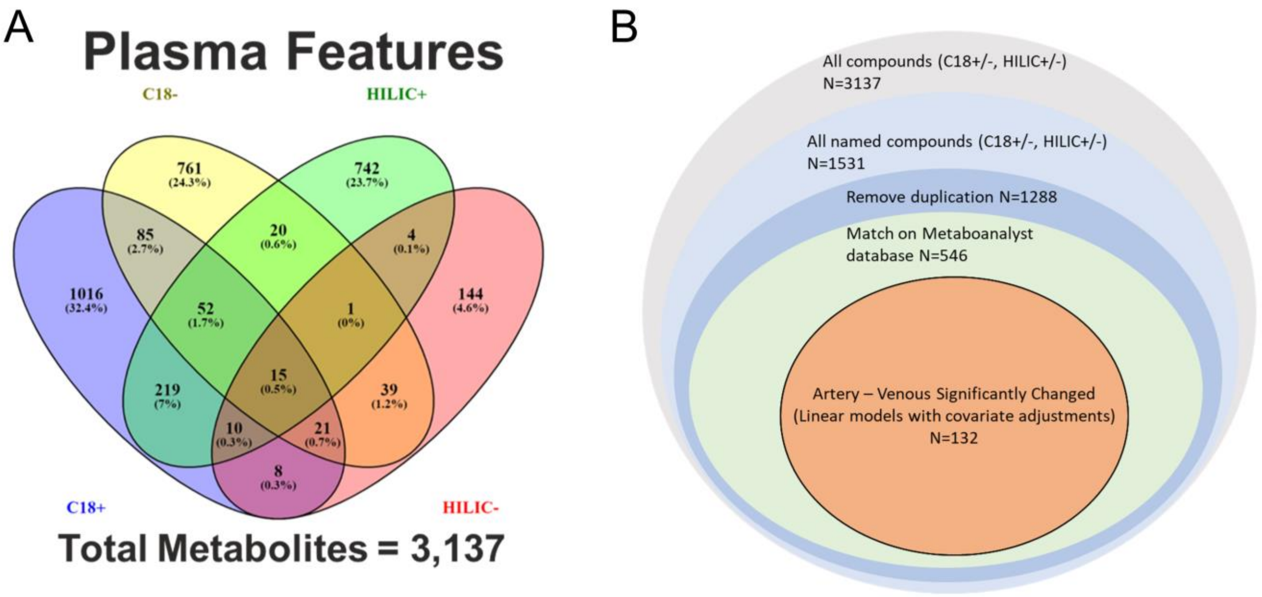

**Figure S12**. Metabolite features in plasma

Venn diagram shows the number of metabolites detected in each of 4 modes (C18+/-, HILIC+/-) in plasma (A). The number of metabolites detected, named and listed in the Metaboanalyst 5.0 database (November 2022) (B). The number of those that artery and venous difference are significantly ($p<0.05$) differed from LS at HS by linear models with covariate adjustment is also shown.

Figure S13.

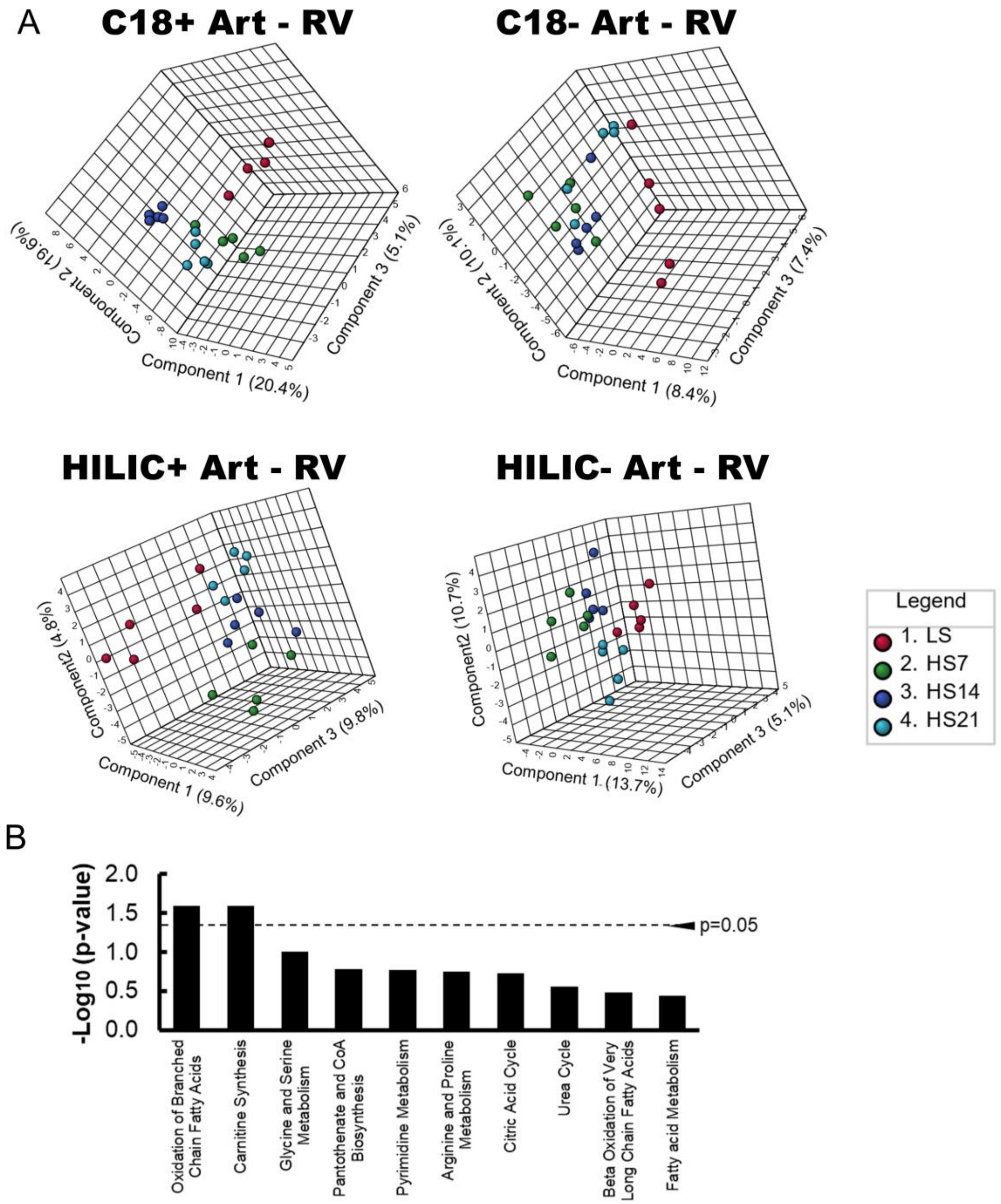

**Figure S13**. sPLS-DA and pathway analysis of arterial and venous difference

(A) Sparse Partial Least-Squares Discriminant Analysis (sPLS-DA) of arterial and venous plasma differences in each of 4 modes (C18+/-, HILIC+/-). Parameters for sPLS-DA are fixed to number of components: 5, variables per component: 20 and validation method: 5-fold CV. Red: Low salt (LS), green: 7 days of high salt (HS), blue: 14 days of HS and light blue: 21 days of HS. (B) Top 10 pathways in order of $-\log_{10}$ (p-value) by enrichment analysis on Metaboanalyst 5.0 (SMPDB, November 2022). Metabolites which are significantly altered by linear models with covariate adjustment are analyzed.

Figure S14.

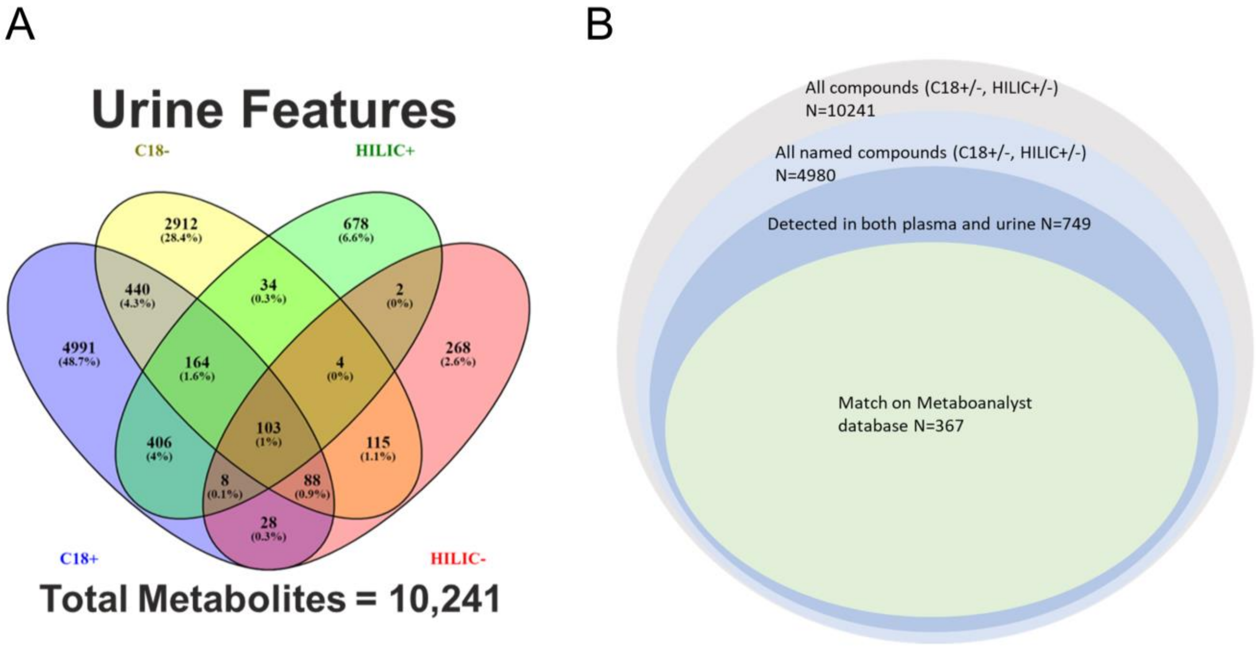

**Figure S14**. Metabolite features in urine

Venn diagram shows the number of metabolites detected in each of 4 modes (C18+/-, HILIC+/-) in urine (A). The number of metabolites detected, named, detected in both urine and plasma, and listed in the Metaboanalyst 5.0 database (November 2022) are also shown(B).

**Table S1**.

| | Age (week) | Body weight (BW; g) | Right kidney weight (RKW; g) | Left kidney weight (LKW; g) | RKW/BW (mg/g) | LKW/BW (mg/g) |
|---|---|---|---|---|---|---|
| LS group | 15.2 ± 0.5 | 360 ± 9 | 1.26 ± 0.02 | 1.23 ± 0.05 | 3.50 ± 0.11 | 3.42 ± 0.12 |
| HS group | 15.3 ± 0.5 | 352 ± 14 | 1.23 ± 0.05 | 1.22 ± 0.05 | 3.50 ± 0.08 | 3.46 ± 0.09 |
| T-test | N.S. | N.S. | N.S. | N.S. | N.S. | N.S. |

**Table S1**. Body and kidney weight data related to rats studied to determine GFR.
Mean ± SEM of the body weights (BW), right kidney weight (RKW), left kidney weight (LKW) and those ratio to BW of a group fed HS for 21 days (n=6) and of rats of the same age fed only LS (n=5). T-test. p>0.05 (N.S.) for all measured parameters.

**Table S2**.

| | Age (week) | GFR (mL/min/100 gbw) | Body weight (g) | Hct (%) | Half life of sinistrin (min) |
|---|---|---|---|---|---|
| LS | 12.3 ± 0.5 | 0.64 ± 0.04 | 305 ± 13 | 39 ± 1 | 33.9 ± 2.0 |
| HS7 | 13.3 ± 0.5 | 0.83 ± 0.05* | 320 ± 12* | 41 ± 1 N.S. | 26.1 ± 1.5* |
| HS14 | 14.3 ± 0.5 | 0.81 ± 0.06* | 339 ± 12* | 41 ± 1 N.S. | 27.1 ± 2.0* |
| HS21 | 15.3 ± 0.5 | 0.85 ± 0.05* | 352 ± 14* | 43 ± 1* | 25.6 ± 1.4* |

**Table S2**. GFR measurements of unanesthetized SD rats studied at LS and at 7, 14, and 21 days of the HS diet.
N=6. Mean ± SEM of age, GFR, body weight, hematocrit (Hct) and half-life of sinistrin.
*p<0.05 vs LS, One-way RM ANOVA, Holm-Sidak. N.S.: p>0.05

**Table S3**.

| | RBF (12h daytime) (mL/min/100 gbw) | RVR (12h daytime) (mmHg/mL/min/100 gbw) | $O_2$ Consumption (mL/min/kgbw) | $O_2$ Delivery (mL/min/kgbw) |
|---|---|---|---|---|
| LS | 3.03 ± 0.18 | 37.2 ± 1.5 | 0.495 ± 0.057 | 4.88 ± 0.22 |
| HS7 | 3.38 ± 0.18* | 35.9 ± 1.8 $^{N.S.}$ | 0.638 ± 0.033* | 5.40 ± 0.33 $^{N.S.}$ |
| HS14 | 3.36 ± 0.17* | 36.2 ± 2.0 $^{N.S.}$ | 0.630 ± 0.061* | 5.31 ± 0.27 $^{N.S.}$ |
| HS21 | 3.33 ± 0.18* | 36.2 ± 1.0 $^{N.S.}$ | 0.741 ± 0.035* | 5.31 ± 0.34 $^{N.S.}$ |

**Table S3**. Renal blood flow (RBF), calculated renal vascular resistance (RVR), $O_2$ consumption, and $O_2$ delivery normalized by body weight of the GFR group of rats.

N=6. Mean ± SEM, *$p<0.05$ vs LS, One-way RM ANOVA, Holm-Sidak.  N.S.: $p>0.05$

**Table S4**.

| Renal Vein | | | | | | | | | |
|---|---|---|---|---|---|---|---|---|---|
| | Hb (g/dL) | pO2 (mmHg) | SHbO$_2$ (%) | O$_2$ content (mL/dL) | pCO2 (mmHg) | Na+ (mM) | K+ (mM) | Cl- (mM) | Ca++ (mM) |
| LS | 13.8 (0.5) | 77.4 (3.0) | 86.4 (1.1) | 15.8 (0.4) | 35.2 (1.6) | 144.8 (0.5) | 3.3 (0.1) | 115.0 (1.3) | 1.3 (0.2) |
| HS7 | 13.8 N.S. (0.4) | 75.8 N.S. (1.1) | 83.8 N.S. (0.9) | 15.4 N.S. (0.5) | 35.7 N.S. (1.0) | 144.7 N.S. (0.7) | 3.4 N.S. (0.1) | 112.3 N.S. (1.3) | 1.1 N.S. (0.0) |
| HS14 | 13.5 N.S. (0.4) | 72.8 N.S. (1.3) | 82.4* (1.1) | 14.8* (0.5) | 36.1 N.S. (2.2) | 144.5 N.S. (1.1) | 3.7* (0.1) | 116.5 N.S. (3.3) | 1.0 N.S. (0.1) |
| HS21 | 13.3 N.S. (0.4) | 69.1 N.S. (2.7) | 81.4* (1.0) | 14.4* (0.6) | 34.5 N.S. (1.2) | 145.5 N.S. (1.1) | 3.6 N.S. (0.1) | 116.2 N.S. (2.8) | 1.0 N.S. (0.0) |

| Artery | | | | | | | | | |
|---|---|---|---|---|---|---|---|---|---|
| | Hb (g/dL) | pO2 (mmHg) | SHbO$_2$ (%) | O$_2$ content (mL/dL) | pCO2 (mmHg) | Na+ (mM) | K+ (mM) | Cl- (mM) | Ca++ (mM) |
| LS | 13.4 (0.2) | 116.8 (2.9) | 95.6 (0.6) | 17.6 (0.6) | 32.0 (2.4) | 146.2 (0.9) | 3.0 (0.1) | 117.8 (2.4) | 1.3 (0.1) |
| HS7 | 13.6 N.S. (0.5) | 110.3* (2.9) | 94.8 N.S. (0.3) | 17.5 N.S. (0.4) | 34.4 N.S. (1.2) | 145.5 N.S. (0.6) | 3.1 N.S. (0.1) | 113.0 N.S. (1.6) | 1.1 N.S. (0.0) |
| HS14 | 14.1 N.S. (0.5) | 104.2* (1.8) | 93.4* (0.3) | 16.9 N.S. (0.5) | 35.4 N.S. (2.0) | 144.7 N.S. (1.3) | 3.3 N.S. (0.1) | 115.2 N.S. (3.9) | 1.2 N.S. (0.1) |
| HS21 | 13.7 N.S. (0.4) | 109.3* (2.9) | 94.1* (0.5) | 16.7 N.S. (0.5) | 33.1 N.S. (1.6) | 145.8 N.S. (1.4) | 3.3 N.S. (0.2) | 114.7 N.S. (3.1) | 1.1 N.S. (0.1) |

**Table S4**. Arterial and renal venous blood gas data.

Total Hemoglobin (Hb), partial pressure of oxygen (pO2), oxyhemoglobin saturation (SHbO2), $O_2$ content and whole blood electrolyte data obtained from the unanesthetized instrumented rats when fed LS, and HS for 7, 14, and 21 days. N=6, Mean and (SEM), *p<0.05 vs LS, One-way RM ANOVA, Holm-Sidak. N.S.: p>0.05 (N.S.), mM: mmol/L

**Table S5**.

| Cortex | | | | | | Outer medulla | | | | | |
|---|---|---|---|---|---|---|---|---|---|---|---|
| Gene name | Log2 FC (HS14/LS) | Adj. p-value LS-HS14 | Gene name | Log2 FC (HS21/LS) | Adj. p-value LS-HS21 | Gene name | Log2 FC (HS14/LS) | Adj. p-value LS-HS14 | Gene name | Log2 FC (HS21/LS) | Adj. p-value LS-HS21 |
| Stc1 | 1.45 | 1.0E-03 | Lyz2 | 2.74 | 8.1E-06 | LOC100911372 | 1.78 | 2.7E-01 | RT1-A2 | 2.15 | 3.6E-03 |
| Chchd10 | 1.37 | 2.6E-01 | C3 | 2.47 | 1.2E-05 | Matn1 | 1.71 | 7.3E-02 | Itgal | 2.13 | 1.2E-14 |
| Matn1 | 1.34 | 5.3E-06 | RT1-A2 | 2.42 | 2.1E-05 | C7 | 1.59 | 5.7E-07 | C3 | 2.02 | 1.3E-07 |
| Grem1 | 1.31 | 2.2E-02 | Itgal | 2.32 | 8.2E-12 | Svep1 | 1.59 | 1.4E-03 | Ptprc | 1.70 | 1.3E-06 |
| Lyz2 | 1.13 | 5.9E-07 | RT1-Db1 | 2.32 | 7.6E-10 | Col12a1 | 1.57 | 5.9E-02 | Tap1 | 1.69 | 1.9E-04 |
| C1s | 1.07 | 9.9E-05 | Fcgr3a | 2.20 | 3.7E-05 | Nr4a1 | 1.55 | 7.9E-01 | C7 | 1.67 | 1.8E-06 |
| Mx2 | 1.06 | 5.6E-01 | Psmb9 | 2.19 | 7.5E-07 | Cp | 1.55 | 3.3E-03 | Lyz2 | 1.60 | 1.8E-02 |
| Spp1 | 1.05 | 3.2E-02 | Gbp2 | 2.17 | 8.3E-05 | Sfrp1 | 1.53 | 4.3E-03 | Irf1 | 1.59 | 1.5E-04 |
| C3 | 1.00 | 1.6E-04 | Trpm2 | 2.11 | 1.2E-07 | C1s | 1.40 | 1.5E-01 | RT1-Da | 1.59 | 9.5E-05 |
| Pkhd1 | 0.97 | 3.2E-02 | Tap1 | 2.07 | 1.8E-05 | Smoc2 | 1.37 | 5.9E-02 | Itgax | 1.47 | 4.6E-17 |
| Rps19 | 0.96 | 3.2E-01 | Rac2 | 1.97 | 1.5E-10 | Lrp1 | 1.37 | 2.8E-02 | Cp | 1.46 | 5.4E-03 |
| Vps13a | 0.95 | 5.5E-02 | Laptm5 | 1.92 | 2.3E-07 | C1r | 1.31 | 1.1E-01 | Coro1a | 1.44 | 1.3E-05 |
| RT1-Db1 | 0.94 | 7.7E-02 | C1qb | 1.90 | 9.9E-05 | Cdh11 | 1.30 | 3.4E-02 | Cybb | 1.43 | 4.5E-04 |
| Dnase1 | 0.94 | 1.2E-08 | Ctsz | 1.87 | 3.6E-12 | Pkm | 1.23 | 4.5E-01 | C1s | 1.43 | 3.2E-02 |
| Plekhh1 | 0.94 | 9.5E-02 | Ptprc | 1.87 | 5.3E-09 | Lamc3 | 1.23 | 2.0E-02 | Cd74 | 1.41 | 1.7E-05 |
| Slc16a7 | 0.93 | 7.7E-02 | C1qa | 1.85 | 1.7E-05 | Abca8a | 1.22 | 1.4E-02 | Fcer1g | 1.40 | 4.6E-04 |
| P2rx4 | 0.93 | 6.9E-02 | Apol11a | 1.82 | 1.1E-09 | Cdkn1c | 1.22 | 1.8E-02 | Akna | 1.40 | 9.3E-04 |
| Prodh | 0.91 | 3.8E-07 | Irf1 | 1.82 | 1.1E-06 | Itgal | 1.21 | 2.4E-02 | Abca8a | 1.34 | 1.3E-03 |
| Itgal | 0.89 | 1.3E-04 | Ncf1 | 1.81 | 2.7E-10 | Igsf10 | 1.20 | 7.1E-02 | C1qc | 1.33 | 2.4E-02 |
| Abca8a | 0.88 | 2.1E-04 | Coro1a | 1.81 | 2.4E-14 | Filip1l | 1.18 | 1.0E-02 | Vcam1 | 1.31 | 1.7E-03 |
| Acaa2 | -0.78 | 2.0E-09 | Slc4a4 | -1.35 | 6.1E-43 | Slc26a1 | -0.92 | 9.8E-01 | AY172581.9 | -0.98 | 3.0E-02 |
| Slc22a12 | -0.79 | 1.9E-03 | Dgkg | -1.37 | 1.3E-20 | Proc | -0.95 | 5.9E-01 | Pklr | -0.99 | 6.7E-01 |
| Gstp1 | -0.79 | 3.6E-04 | Esr1 | -1.37 | 1.2E-20 | Glyatl2 | -0.98 | 9.1E-01 | Lifr | -1.03 | 6.4E-01 |
| Tmem37 | -0.81 | 5.5E-11 | Apcs | -1.40 | 2.6E-18 | Entpd2 | -0.99 | 9.8E-07 | Mtr | -1.04 | 2.0E-01 |
| Cyp4a2 | -0.82 | 6.5E-08 | Slc5a8 | -1.41 | 5.6E-14 | Nkd2 | -0.99 | 8.6E-01 | Rpl7a | -1.08 | 2.8E-01 |
| Gjb2 | -0.84 | 1.4E-16 | Ugt2b35 | -1.41 | 9.9E-07 | Aldh1b1 | -1.00 | 8.6E-01 | Cyr61 | -1.10 | 6.5E-02 |
| Tmem86a | -0.87 | 2.5E-02 | Gatm | -1.49 | 2.7E-04 | Ubash3b | -1.00 | 1.7E-01 | LOC100911238 | -1.11 | 1.7E-01 |
| Mki67 | -0.89 | 2.2E-03 | AABR07015081.2 | -1.50 | 6.3E-02 | Eri2 | -1.01 | 9.3E-01 | Nkd2 | -1.12 | 4.6E-01 |
| Cyp4a2 | -0.90 | 6.6E-08 | Epha4 | -1.51 | 4.1E-33 | Ppic | -1.05 | 9.8E-01 | RGD1563354 | -1.14 | 2.9E-02 |
| Pex11a | -0.91 | 2.3E-09 | Arfgef3 | -1.53 | 5.0E-04 | Slc22a12 | -1.06 | 8.6E-01 | Ihh | -1.23 | 1.3E-01 |
| Mlph | -1.05 | 5.6E-07 | Papln | -1.56 | 3.6E-25 | Gpx3 | -1.08 | 9.5E-01 | AY172581.24 | -1.29 | 3.2E-02 |
| Papln | -1.08 | 1.7E-10 | Tgm2 | -1.58 | 2.3E-12 | Slc17a3 | -1.13 | 9.4E-01 | Dusp1 | -1.32 | 2.6E-02 |
| Eci1 | -1.09 | 1.7E-10 | Hnrnpab | -1.60 | 1.6E-01 | RGD1563354 | -1.19 | 1.4E-01 | Nr4a1 | -1.32 | 6.9E-02 |
| Ugt2b35 | -1.10 | 2.0E-03 | Vnn1 | -1.65 | 1.4E-14 | Rimbp2 | -1.27 | 6.1E-01 | LOC100910882 | -1.42 | 8.8E-02 |
| Tgm2 | -1.15 | 1.7E-02 | Mlph | -1.96 | 1.7E-15 | Pklr | -1.27 | 5.3E-01 | Junb | -1.43 | 3.0E-02 |
| Mthfr | -1.23 | 9.8E-04 | Slc16a1 | -2.16 | 4.8E-13 | AABR07015081.2 | -1.29 | 5.2E-01 | AABR07015081.1 | -2.06 | 8.1E-04 |
| Vnn1 | -1.35 | 3.9E-09 | Nrep | -2.24 | 8.3E-32 | Slc22a6 | -1.41 | 8.5E-01 | AABR07063425.2 | -2.14 | 5.1E-04 |
| Nrep | -1.47 | 1.4E-16 | Mthfr | -2.27 | 2.0E-14 | Acsm3 | -1.45 | 8.8E-01 | AABR07015057.1 | -2.18 | 2.4E-04 |
| Prima1 | -2.39 | 9.4E-11 | Prima1 | -2.35 | 3.3E-35 | LOC100910882 | -1.75 | 1.5E-01 | AABR07015081.2 | -2.32 | 4.7E-06 |
| Hmgcs2 | -2.73 | 1.8E-10 | Hmgcs2 | -3.00 | 2.9E-18 | AABR07063425.2 | -1.87 | 9.9E-02 | Egr1 | -3.32 | 8.6E-05 |

**Table S5**. Top and bottom 20 genes with large fold change (FC) between high salt (HS) and low salt (LS) group in cortex (Cx) and outer medulla (OM).

**Table S6**.

| | | |
|---|---|---|
| (2E,4E)-2,4-Dodecadienal | Butenylcarnitine | L-Norleucine |
| (3β,5α,9α,22E,24R)-3,5,9-Trihydroxy-23-methylergosta-7,22-dien-6-one | Bz-Arg-OEt | L-Palmitoylcarnitine |
| (3E)-4-(4-hydroxyphenyl)but-3-en-2-one | Capsiamide | L-Tryptophan |
| 1,11-Undecanedicarboxylic acid | Cepanone | L-Tyrosine |
| 1,2-Benzisothiazol-3(2H)-one | Cholic acid | L-Valine |
| 1,2-Dihydro-1,1,6-trimethylnaphthalene | Citric acid | LysoPC(22:5(7Z,10Z,13Z,16Z,19Z)) |
| 1-Methylguanine | Citrulline | Methylphosphate |
| 2,2-Bis[4-(2,3-epoxypropoxy)phenyl]propane | Creatine | N-(5-Methyl-3-oxohexyl)alanine |
| 2,4,6-Octatriyn-1-ol | Creatinine | Nalidixic Acid |
| 2,5-Furandicarboxylic acid | Cyclohexylamine | N-Desmethyltramadol |
| 2-acetyl-1-alkyl-sn-glycero-3-phosphocholine | Cytidine | N-methyl-L-glutamic Acid |
| 2-Acetyl-4-methylpyridine | Cytidine monophosphate | Norophthalmic acid |
| 2-Amino-9,10-epoxy-8-oxodecanoic acid | Decarbamoylsaxitoxin | N-Undecanoylglycine |
| 2-Diethylaminoethanol | Deoxycytidine | Ornithine |
| 2-Ethylglutaric acid | Dibutyl phthalate | o-Xylene |
| 2-Methoxyestrone | Diethylene glycol | Pantothenic acid |
| 3,3-Dimethylglutaric acid | Dihydrothymine | p-Cresol sulfate |
| 3,4-Dihydroxyhydrocinnamic acid | Dodecanoic acid | Pelargonic acid |
| 3,4-Methylenesebacic acid | Dodecenoic acid | Pentylbenzene |
| 3-Buten-1-amine | D-Pipecolic acid | Phenylacetylglycine |
| 3-Hydroxycapric acid | D-Proline | Phthalic acid |
| 3-Hydroxytetradecanedioic acid | Eicosapentaenoic acid | Propionylcarnitine |
| 3-hydroxytridecanoic acid | Epigallocatechin | Prostaglandin B1 |
| 3-Oxooctanoic acid | Ethyl lactate | Pyridoxamine |
| 4-Acetamidobenzoic acid | Etodolac | Pyrrolidonecarboxylic acid |
| 4-Hydroxy-3-methoxybenzenemethanol | gamma-Asarone | Pyruvic acid |
| 4-Hydroxybenzaldehyde | Glycocholic acid | Salicylic acid |
| 4-Hydroxybutyric acid | Hexanoylcarnitine | Sebacic acid |
| 4-Trimethylammoniobutanoic acid | Hexylbenzene | Tartaric acid |
| 5-Hydroxy-2-furoic acid | Hydroxyphenyllactic acid | Taurochenodesoxycholic acid |
| 5-hydroxy-2-oxo-4-ureido-2,5-dihydro-1H-imidazole-5-carboxylate | Indole-3-methyl acetate | Tetraethylene glycol |
| 5-Methylcytidine | Indoleacetic acid | Thiamine |
| 5-Methylcytosine | Isovaleric acid | Thiorphan |
| 7-Oxoheptanoic acid | L-Acetylcarnitine | trans-Aconitic acid |
| 8-Amino-7-oxononanoic acid | L-Carnitine | Traumatic acid |
| 9-cis-Retinoic acid | L-Dihydroanticapsin | Tridecanoic acid |
| Adipic acid | L-Glutamine | Triethylamine |
| Allantoin | L-Histidinal | Trimethadione |
| Aspirin | L-Homoserine | Trimethylamine N-oxide |
| Asymmetric dimethylarginine | L-Isoleucine | Ureidopropionic acid |
| Benzaldehyde | L-Kynurenine | Uric acid |
| Benzoic acid | L-Lactic acid | Valproic acid |
| Betaine | L-Lysine | Violet-leaf aldehyde |
| Betonicine | L-Methionine | α-Ketoglutaric acid |

**Table S6**. A-V significantly changed metabolites over time by linear models with covariate adjustments.

All of the n=131 metabolites which are matched on Metaboanalyst 5.0 database (November 2022) and whose arterial and venous plasma differences are significantly ($p<0.05$) changed by high salt (HS) over time by linear models with covariate adjustments.

**Table S7**.

| Metabolites detected in both plasma and urine N=367 | | | |
|---|---|---|---|
| Metabolites excreted in urine in excess of filtration fraction | | | |
| LS (N=19) | HS7 (N=29) | HS14 (N=37) | HS21 (N=42) |
| (S)-Pinocembrin | (S)-Pinocembrin | (S)-Pinocembrin | (S)-Pinocembrin |
| 2-Diethylaminoethanol | 2-Diethylaminoethanol | 2-Diethylaminoethanol | 2-Diethylaminoethanol |
| 2-Hydroxystearic acid | 2-Hydroxystearic acid | 2-Hydroxystearic acid | 2-Hydroxystearic acid |
| Biotin | Biotin | Biotin | Biotin |
| Capryloylglycine | Capryloylglycine | Capryloylglycine | Capryloylglycine |
| D-Glyceraldehyde 3-phosphate | D-Glyceraldehyde 3-phosphate | D-Glyceraldehyde 3-phosphate | D-Glyceraldehyde 3-phosphate |
| Kynurenic acid | Kynurenic acid | Kynurenic acid | Kynurenic acid |
| L-Methionine | L-Methionine | L-Methionine | L-Methionine |
| N-Acetylvaline | N-Acetylvaline | N-Acetylvaline | N-Acetylvaline |
| Nalidixic Acid | Nalidixic Acid | Nalidixic Acid | Nalidixic Acid |
| Phenylacetylglycine | Phenylacetylglycine | Phenylacetylglycine | Phenylacetylglycine |
| Porphobilinogen | Porphobilinogen | Porphobilinogen | Porphobilinogen |
| Pseudouridine | Pseudouridine | Pseudouridine | Pseudouridine |
| Triacetin | Triacetin | Triacetin | Triacetin |
| Tyramine | Tyramine | Tyramine | Tyramine |
| Uric acid | Uric acid | Uric acid | Uric acid |
| Propionylcarnitine | Propionylcarnitine | Propionylcarnitine | |
| Methylsuccinic acid | | | |
| Tiglylcarnitine | | | |
| | 3-Hydroxyanthranilic acid | 3-Hydroxyanthranilic acid | 3-Hydroxyanthranilic acid |
| | Alanyl-Isoleucine | Alanyl-Isoleucine | Alanyl-Isoleucine |
| | Creatinine | Creatinine | Creatinine |
| | Cyclohexylamine | Cyclohexylamine | Cyclohexylamine |
| | Guaifenesin | Guaifenesin | Guaifenesin |
| | Hippuric acid | Hippuric acid | Hippuric acid |
| | Indoxyl sulfate | Indoxyl sulfate | Indoxyl sulfate |
| | L-Dihydroanticapsin | L-Dihydroanticapsin | L-Dihydroanticapsin |
| | N-Acetylarylamine | N-Acetylarylamine | N-Acetylarylamine |
| | Suberic acid | Suberic acid | Suberic acid |
| | Bethanidine | | |
| | Hexanal | | |
| | | 17a-Ethynylestradiol | 17a-Ethynylestradiol |
| | | 4-Oxoproline | 4-Oxoproline |
| | | Dethiobiotin | Dethiobiotin |
| | | Indole | Indole |
| | | Methylglutaric acid | Methylglutaric acid |
| | | Oxoglutaric acid | Oxoglutaric acid |
| | | Uracil | Uracil |
| | | 4-Trimethylammoniobutanoic acid | 4-Hydroxy-3-methoxybenzenemethanol |
| | | Glutarylcarnitine | Allantoin |
| | | Pantothenic acid | Aromadendrin |
| | | | Hypoxanthine |
| | | | Inosine |
| | | | Methylsuccinic acid |
| | | | p-Cresol sulfate |
| | | | Salicylic acid |
| | | | Serotonin |

**Table S7**. Urinary metabolism whose excretion is higher than filtration fraction

The number of metabolites detected in both plasma and urine, and that excreted in excess of filtration fraction are shown.

All of the metabolites which excreted in excess of filtration fraction are listed.

**Table S8**.

| | Artery (Art) (mM) | Renal vein (RV) (mM) | Art-RV (mM) |
|---|---|---|---|
| LS | 1.6 ± 0.8 | 1.3 ± 0.3 | 0.4 ± 0.6 |
| HS7 | 0.9 ± 0.2 N.S. | 1.0 ± 0.1N.S. | -0.1 ± 0.1 N.S. |
| HS14 | 0.7 ± 0.1 N.S. | 0.9 ± 0.1 N.S. | -0.1 ± 0.1 N.S. |
| HS21 | 0.8 ± 0.1 N.S | 1.1 ± 0.1 N.S. | -0.2 ± 0.1 N.S. |

**Table S8**. Validation of lactate concentration in arterial and renal venous plasma by fluorometric assay kit. Mean ± SEM and individual data. One-way ANOVA, No significant difference (p>0.05, n.s.) between groups. mM: mmol/L
