## Supplementary material for "Metabolic responses of normal rat kidneys to a high salt intake": mRNAseq method

### Sample Quality Control

Please refer to QC report for methods of sample quality control.

### Library preparation for Transcriptome sequencing

Messenger RNA was purified from total RNA using poly-T oligo-attached magnetic beads. After fragmentation, the first strand cDNA was synthesized using random hexamer primers, followed by the second strand cDNA synthesis using either dUTP for directional library or dTTP for non-directional library.

For the non-directional library, it was ready after end repair, A-tailing, adapter ligation, size selection, amplification, and purification(**Figure A**).

For the **directional** library, it was ready after end repair, A-tailing, adapter ligation, size selection, **USER enzyme digestion**, amplification, and purification(**Figure B**).

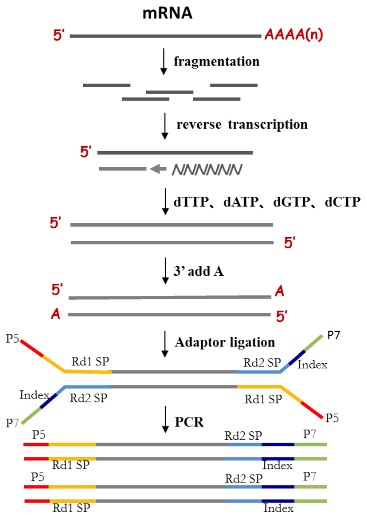


Figure A Workflow of non-directional library construction

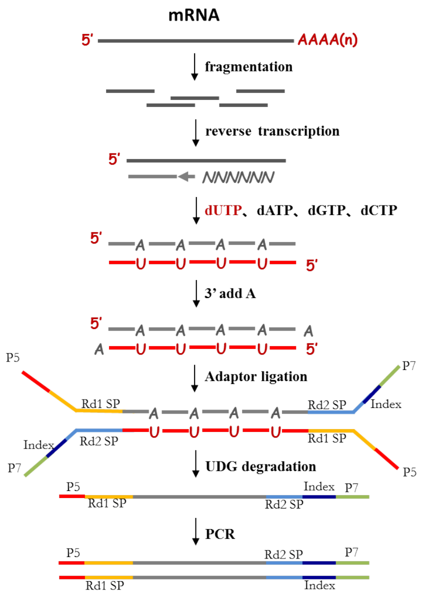


Figure B Workflow of directional library construction

The library was checked with Qubit and real-time PCR for quantification and bioanalyzer for size distribution detection. Quantified libraries will be pooled and sequenced on Illumina platforms, according to effective library concentration and data amount.

#### Clustering and sequencing (Novogene Experimental Department)

The clustering of the index-coded samples was performed according to the manufacturer's instructions. After cluster generation, the library preparations were sequenced on an Illumina platform and paired-end reads were generated.

### Data Analysis

#### Quality control

Raw data (raw reads) of fastq format were firstly processed through in-house perl scripts. In this step, clean data (clean reads) were obtained by removing reads containing adapter, reads containing ploy-N and low quality reads from raw data. At the same time, Q20, Q30 and GC content the clean data were calculated. All the downstream analyses were based on the clean data with high quality.

#### Reads mapping to the reference genome

Reference genome and gene model annotation files were downloaded from genome website directly. Index of the reference genome was built using Hisat2 v2.0.5 and paired-end clean reads were aligned to the reference genome using Hisat2 v2.0.5. We selected Hisat2 as the mapping tool for that Hisat2 can generate a database of splice junctions based on the gene model annotation file and thus a better mapping result than other non-splice mapping tools.

**Novel transcripts prediction**

The mapped reads of each sample were assembled by StringTie (v1.3.3b) (Mihaela Pertea.et al. 2015) in a reference-based approach. StringTie uses a novel network flow algorithm as well as an optional de novo assembly step to assemble and quantitate fulllength transcripts representing multiple splice variants for each gene locus.

**Quantification of gene expression level**

featureCounts v1.5.0-p3 was used to count the reads numbers mapped to each gene. And then FPKM of each gene was calculated based on the length of the gene and reads count mapped to this gene. FPKM, expected number of Fragments Per Kilobase of transcript sequence per Millions base pairs sequenced, considers the effect of sequencing depth and gene length for the reads count at the same time, and is currently the most commonly used method for estimating gene expression levels.

**Differential expression analysis**

*(For DESeq2 with biological replicates)* Differential expression analysis of two conditions/groups (two biological replicates per condition) was performed using the DESeq2 R package (1.20.0). DESeq2 provide statistical routines for determining differential expression in digital gene expression data using a model based on the negative binomial distribution. The resulting P-values were adjusted using the Benjamini and Hochberg's approach for controlling the false discovery rate . Genes with an adjusted P-value <=0.05 found by DESeq2 were assigned as differentially expressed.

*(For edgeR without biological replicates)* Prior to differential gene expression analysis, for each sequenced library, the read counts were adjusted by edgeR program package through one scaling normalized factor. Differential expression analysis of two conditions was performed using the edgeR R package (3.22.5). The P values were adjusted using the Benjamini & Hochberg method. Corrected P-value of 0.05 and absolute foldchange of 2 were set as the

threshold for significantly differential expression.

**GO and KEGG enrichment analysis of differentially expressed genes**

Gene Ontology (GO) enrichment analysis of differentially expressed genes was implemented by the clusterProfiler R package, in which gene length bias wascorrected. GO terms with corrected Pvalue less than 0.05 were considered significantly enriched by differential expressed genes. KEGG is a database resource for understanding high-level functions and utilities of the biological system, such as the cell, the organism and the ecosystem, from molecular-level information, especially large-scale molecular datasets generated by genome sequencing and other high-through put experimental technologies (http://www.genome.jp/kegg/). We used clusterProfiler R package to test the statistical enrichment of differential expression genes in KEGG pathways.

**Gene Set Enrichment Analysis**

Gene Set Enrichment Analysis (GSEA) is a computational approach to determine if a pre-defined Gene Set can show a significant consistent difference between two biological states. The genes were ranked according to the degree of differential expression in the two samples, and then the predefined Gene Set were tested to see if they were enriched at the top or bottom of the list. Gene set enrichment analysis can include subtle expression changes. We use the local version of the GSEA analysis tool http://www.broadinstitute.org/gsea/index.jsp, GO, KEGG data set were used for GSEA independently.

**SNP analysis**

GATK (v4.1.1.0) software was used to perform SNP calling. Raw vcf files were filtered with GATK standard filter method and other parameters (cluster:3; WindowSize:35; QD < 2.0 ; FS > 30.0; DP < 10.

**AS analysis**

Alternative Splicing is an important mechanism for regulate the expression of genes and the variable of protein. rMATS(4.1.0) software was used to analysis the AS event.

**PPI analysis of differentially expressed genes**

PPI analysis of differentially expressed genes was based on the STRING database, which known and predicted Protein-Protein Interactions.
